# P2X4 Purinergic Receptors as a Therapeutic Target in Aggressive Prostate Cancer

**DOI:** 10.1101/2021.06.04.446195

**Authors:** Janielle P. Maynard, Jiayun Lu, Igor Vidal, Jessica Hicks, Luke Mummert, Tamirat Ali, Ryan Kempski, Ayanna M. Carter, Rebecca Sosa, Lauren B. Peiffer, Corinne E. Joshu, Tamara L. Lotan, Angelo M. De Marzo, Karen S. Sfanos

## Abstract

Prostate cancer (PCa) remains a leading cause of cancer-related deaths in American men and treatment options for metastatic PCa are limited. There is a critical need to identify new mechanisms that contribute to PCa progression, that distinguish benign from lethal disease, and that have potential for therapeutic targeting. P2X4 belongs to the P2 purinergic receptor family that is commonly upregulated in cancer and is associated with poorer outcomes. Herein, we report that the P2X4 purinergic receptor is overexpressed in PCa, associated with PCa metastasis, and a driver of tumor development *in vivo*. We observed P2X4 protein expression primarily in epithelial cells of the prostate, a subset of CD66^+^ neutrophils, and most CD68^+^ macrophages. Our analysis of tissue microarrays representing 491 PCa cases demonstrated significantly elevated P2X4 expression in cancer compared to benign tissue spots, in prostatic intraepithelial neoplasia, in cancer from White compared to Black men, and in PCa with ERG positivity or with PTEN loss. High P2X4 expression in benign tissues was likewise associated with the development of metastasis after radical prostatectomy. Treatment with P2X4-specific agonist CTP increased transwell migration and invasion of PC3, DU145, and CWR22Rv1 PCa cells. P2X4 antagonist 5-BDBD treatment resulted in a dose-dependent decrease in viability of PC3, DU145, LNCaP, CWR22Rv1, TRAMP-C2, Myc-CaP, BMPC1, and BMPC2 cells and decreased DU145 cell migration and invasion. Knockdown of P2X4 attenuated growth, migration, and invasion of PCa cells. Finally, knockdown of P2X4 in Myc-CaP cells resulted in significantly attenuated subcutaneous allograft growth in FVB/NJ mice. Collectively, these data strongly support a role for the P2X4 purinergic receptor in PCa aggressiveness and identifies P2X4 as a candidate for therapeutic targeting.

## Introduction

Prostate cancer (PCa) remains the second leading cause of cancer-related deaths in American men and Black men are two to three times more likely to die from the disease than White men are (1,2). Currently, we lack effective therapies against distant metastatic PCa for which the 5-year survival is about 30% (1). There is a critical need to identify new mechanisms that contribute to PCa progression, that distinguish benign from lethal disease, and that have potential for therapeutic targeting.

Chronic inflammation may contribute to prostate carcinogenesis and is associated with aggressive prostate cancer (3). Exogenous exposures stimulate innate immune cells including macrophages and neutrophils to infiltrate the prostate. These phagocytes release reactive oxygen species and reactive nitrogen species causing DNA damage, cell injury, and cell death (4,5). Cell stress, cell injury, and cell death subsequently result in the release of danger-associated molecular patterns (DAMPs), including extracellular ATP (eATP), that exacerbate the inflammatory response (6,7). Increased eATP concentrations act as powerful chemotactic stimuli for host innate immune cells, many of which express P2 purinergic receptors (6,8-10). P2X purinergic receptors are ionotropic ligand-gated ion channel-type receptors (P2X1-P2X7) and P2Y purinergic receptors are metabotropic G protein-coupled receptors (P2Y1, P2Y2, P2Y4, P2Y6, P2Y11-14) (11,12). P2 purinergic receptors are differentially expressed across cell types and may sometimes only be detected in pathological conditions (11). When eATP binds P2 purinergic receptors on immune cells, activation results in the induction of chemokines and inflammatory markers, and the assembly or activation of the inflammasome (13-15). These events help drive inflammation and further stimulate ATP release from nearby cells. Propagation of inflammation is thought, in part, to upregulate multiple P2 purinergic receptors on both immune and non-immune cells (16). Activation of P2 purinergic receptors on non-immune cells promotes cell proliferation and mediates cell differentiation promoting migratory and invasive phenotypes (17-19). For instance, P2X7 and P2Y2 activation reportedly improves the invasiveness of PCa cells (20,21). Consequently, purinergic signaling has been studied in the context of inflammation and cancer. Significantly greater eATP concentrations were measured in models of chronic inflammation and in the tumor interstitium of mice compared to healthy tissues (22,23). P2 purinergic receptor overexpression is reported in multiple cancer types and correlates with poorer outcomes (24-28). P2X4 purinergic receptor expression was identified as the most highly expressed P2 purinergic receptor in PC3, LNCaP, and C4-2B PCa cell lines and P2X4 antagonists had anti-tumorigenic effects in a PCa xenograft model (29). However, the cellular localization and function of P2 purinergic receptors and their association with PCa outcomes is not fully described.

Herein, we demonstrate elevated P2X4 purinergic receptor expression in prostatic intraepithelial neoplasia (PIN) and PCa compared to benign tissues. P2X4 receptor expression was significantly increased in cases with ERG positivity or with PTEN loss and higher P2X4 protein expression in benign regions adjacent to primary PCa was associated with an increased risk of the development of metastasis after radical prostatectomy. We further demonstrate that P2X4 receptor drives PCa cell growth, migration, and invasion *in vitro* and PCa tumor development *in vivo*.

## Materials and Methods

### Patient population and clinical samples

All specimens were acquired using protocols approved by the Johns Hopkins University Institutional Review Board. Formalin-fixed paraffin-embedded (FFPE) whole tissue slides (e.g. standard tissue slides) from tissue blocks were obtained from organ donor prostates with no indication of cancer from 1 self-identified Black and 3 self-identified White men. Tissue slides from tumor containing block, matched benign tissue block with inflammation, and a matched benign tissue block without overt inflammation from previously untreated patients with clinically localized PCa were obtained from radical prostatectomy specimens from 3 self-identified Black patients and 3 self-identified White patients (n = 6 patients with whole tissue slides) with lower-grade (Gleason sum ≤ 3+4, ISUP Grade Group 1-2) PCa. Three separate tissue microarray (TMA) sets were used in this study: Prostate Cancer Biorepository Network (PCBN) High Grade Race, Race Disparity (30,31), and Metastasis at Autopsy. The “PCBN High Grade Race” TMA set contains radical prostatectomy tumor and benign tissue from 120 Black men matched 2:1 to 60 White men on age +/-3 years, grade and stage and is enriched for cases with Gleason score ≥ 8. This TMA contains 4 tissue spots of the index tumor from each case as well as 4 benign tissue spots taken from a block containing only normal tissue, with deliberate avoidance of areas with overt inflammation. The “Race Disparity” TMA set contains tumor (4 spots) and benign (3 spots) tissue from a radical prostatectomy cohort of Black men matched to White men selected via stratified random sampling among Gleason score groupings (3+3, 3+4, 4+3, 8, 9-10) as previously described (30,31); 139 Black men, 151 White men and 1 man listed as Other’ were analyzed on the Race Disparity TMA set. Finally, the “Metastasis at Rapid Autopsy” TMA contains tissues from 11 autopsy metastatic sites from 5 Black and 16 White men. TMAs were used in RNA *in situ* hybridization (RISH) and immunohistochemistry (IHC) experiments. Further details of the patients and TMAs are provided in Suppl. Tables S1 and S2.

### Cell lines and cell treatment

TRAMPC2, Myc-CaP, LNCaP, CWR22Rv1, VCaP, and HEK293T cells were obtained from the American Type Culture Collection (ATCC). PC3 and DU145 were obtained from the NCI-Frederick. LAPC4, and MDA PCa 2b, cells were obtained from J.T. Isaacs (Johns Hopkins University). BMPC1, BMPC2, cells were obtained from Angelo De Marzo (Johns Hopkins University) (32). All human cell lines used were authenticated at the Johns Hopkins Genetic Resources Core Facility via short tandem repeat profiling of 9 genomic loci with the GenePrint 10 System (Promega, Madison, WI) before use. Mouse cell lines used were authenticated by ATCC (Manassas, VA) using ABI Prism® 3500xl Genetic Analyzer and data were analyzed using GeneMapper® ID-X v1.2 software (Applied Biosystems). LNCaP, CWR22Rv1, PC3, and DU145 were maintained in RPMI 1640 media containing L-Glutamine and 10% heat-inactivated fetal bovine serum (FBS) at 37°C and 5% CO_2_. LAPC4 cells were maintained in Iscove’s modified Dulbeco’s Medium (IMDM) media containing L-Glutamine, 25mM HEPES, synthetic steroid R1881 and 10% FBS. BMPC1, BMPC2, Myc-CaP, and HEK293T cells were maintained in Dulbecco’s Modified Eagle Medium (DMEM) media containing L-Glutamine and 10% heat-inactivated FBS. TRAMP-C2 cells were cultured in high glucose DMEM (4.5 g/L glucose, ATCC) with 5% heat inactivated FBS, 5% Nu-Serum IV (Corning, Fisher Scientific, Waltham, MA), and 5μg/ml insulin and 10^−8^ M dihydrotestosterone (Corning, Fisher Scientific, Waltham, MA). Cells were treated with 100μM ATP (A2383, Sigma-Aldrich, St. Louis, MO), 100μM or 400μM CTP (Cat. No. C1506, Sigma-Aldrich, St. Louis, MO), or 5-BDBD (Cat. No. SML0450, Sigma-Aldrich, St. Louis, MO) for 24h.

### P2X4 siRNA, shRNA, and CRISPR-cas9 knockdown

Human P2X4 siRNA (Cat. No. 42569, Santa Cruz, Dallas, TX) or control siRNA (Cat. No. 37007, Santa Cruz, Dallas, TX) was transfected into PC3 cells using Lipofectamine (Cat. No. 11668-030, ThermoScientific, Waltham, MA) and total protein extracts were collected after 72hrs.

P2X4 targeted short hairpin RNA (shRNA) human and mouse plasmids were obtained from the JHU ChemCORE. Human P2X4 shRNAs were incorporated into PC3 cells and mouse P2X4 shRNAs we incorporated in Myc-CaP cells by lentiviral transduction and puromycin (1ug/mL) selection done for at least 5 days. Cells were collected as total protein extracts or into FFPE plugs as previously described (33).

Target P2X4 sequences for CRISPR were identified from Human Brunello Library and the Mouse Brie Library (34). Sequences were modified according to the Zhang Lab Target Guide Sequence Cloning Protocol using the lentiCRISPRv2 plasmid (Addgene, Watertown, MA) (34-36). Appropriate gRNA sequence insertion was verified by Sanger Sequencing. LentiCRISPRv2-P2X4gRNA constructs were packaged into virus using HEK293T cells, psPAX2, and pMD2.G (Addgene, Watertown, MA). Virus containing supernatant was filtered using 0.45 μm filter and stored at -20°C. Human P2X4 constructs were transduced into LNCaP and DU145 cells, and mouse P2X4 constructs were transduced into Myc-CaP cells using polybrene (Millipore, Burlington, MA) followed by puromycin selection. P2X4 knockdown was confirmed by Western blot. Cells were collected into FFPE plugs as previously described (33).

### Western Blotting

Total protein extracts were obtained by homogenizing cells in total lysis buffer (25 mM Tris-HCl, pH 7.5, 150mM NaCl, 1.0% Triton X-100, 0.5% sodium deoxycholate, 0.1% sodium dodecyl sulfate and cOmplete mini protease inhibitor cocktail (Cat. No. 11836153001, Sigma-Aldrich, St. Louis, MO) and centrifuging at 12,000 rpm for 20 min (4°C). Equal amounts of total proteins as determined by BCA Protein Assay were analyzed by SDS-polyacrylamide gel electrophoresis with antibodies specific to IL-8 (Cat. No. 18672, Abcam, Cambridge, MA) and prostate specific antigen (Cat. No. 2475, Cell Signaling, Danvers, MA). The protein bands were detected by enhanced chemiluminescent reaction according to the manufacturer’s instructions (ThermoFisher). Blots were probed with antibody specific for human P2X4 (Cat. No. HPA039494, Sigma-Aldrich, St. Louis, MO), mouse P2X4 (Cat. No. APR-002, Alomone Labs, Jerusalem, Israel), E-cadherin (Cat. No. 3195, Cell Signaling, Danvers, MA.), Vimentin (Cat. No. 5741, Cell Signaling, Danvers, MA), and cleaved caspase-3 (Cat. No. 9664, Cell Signaling, Danvers, MA). β-Actin (Cat. No. A5441, Sigma-Aldrich, St. Louis, MO) or TATA binding protein (Cat. No. 51841, Abcam, Cambridge, MA) were used to ensure equal loading of proteins in each lane.

### MTT assay

Cell viability was determined using the 3-(4, 5-dimethylthiazol-2-yl)-2,5-diphenyltetrazolium assay (MTT assay, Cat. No. M5655, Sigma-Aldrich, St. Louis, MO). Cells were plated in 96-well plates, maintained in RPMI media supplemented with 10% heat-inactivated FBS for 24h and treated with 5-BDBD. Cell were exposed to MTT solution (0.7 mg / ml) and incubated at 37 °C for 2 h. The media was removed and 200 μl of dimethyl sulphoxide (DMSO) was added to each well. After shaking the plates for 30 min, the absorbance at 570 nM was measured (background subtraction at 650 nM).

### Transwell Assay

Cells were plated at a known density in the upper chamber of 8.0μM membrane transwells (Cat. No. 3422, Corning, Tewksbury, MA) with or without matrigel coating (Cat. No. 356234, Corning, Tewksbury, MA) in media containing 1% FBS. Transwells were placed in wells with media containing 10% FBS and 5ug/mL fibronectin (Cat. No. f2006, Sigma-Aldrich, St. Louis, MO). Cells were fixed onto the transwell membrane in 10% formalin with 0.01% Triton X-100 for 10 minutes and stained with DAPI. Images were obtained using a fluorescent microscope, 5-10 fields per membrane, and cells counted using ImageJ.

### RISH

RISH was performed using the RNAscope 2.5 FFPE Brown Reagent Kit (Cat. No. 322310, Advanced Cell Diagnostics, Newark, CA) per the manufacturer instructions. Briefly, FFPE tissues were baked at 60 °C for 30 minutes followed by deparaffinization in three changes of 100% xylene for 10 minutes each and two changes of 100% alcohol for 1 minute each. Next, the slides were treated with hydrogen peroxide for 10 minutes at room temperature (RT). The slides were then added to boiling buffer for 15 minutes in a steamer and then treated with protease digestion buffer for 30 minutes at 40 °C. The slides were incubated with a custom RNAscope target probe designed against human P2X4 mRNA (probe region 406 - 1349, NCBI seq. #NM_001256796.1), human CD68 mRNA (probe region 367-1149, NCBI seq. #NM_001040059.1), human MYC mRNA (probe region 536 – 1995, NCBI seq. #NM_002467.4), and human peptidyl prolyl isomerase B (PPIB, probe region 139-989, NCBI seq. #NM_000942.4), also known as cyclophilin B or mouse PPIB (probe region 98 - 856, NCBI seq. #NM_011149.2) as a positive control mRNA for 2 hours at 40°C, followed by signal amplification. 3,3’-diaminobenzidine (DAB) was used for colorimetric detection for 10 minutes at RT. Slides were counterstained with Gill’s Hematoxylin. All FFPE blocks and sectioned slides were maintained at -20 °C soon after collection, to avoid effects of block age on RISH analyses (37). Slides were scanned using a Hamamatsu NanoZoomer-XR. Slides were viewed, images captured, and TMA images segmented using Proscia Concentriq. Image analysis was done using TMAJ and FrIDA software (version 3.18.0) as previously described (37).

### IHC

IHC was performed using the Power Vision+ Poly-HRP IHC kit (Cat. No. PV6119, Leica Biosystems, Wetzlar, Germany). Slides were steamed for 25 minutes in antigen retrieval solution (Cat. No., H-3300, Vector Laboratories, INC. Burlingame, CA) and incubated with rabbit polyclonal anti-human P2X4 (Cat. No. HPA039494, Sigma-Aldrich, St. Louis, MO), rabbit polyclonal anti-mouse P2X4 (Cat. No. APR-002, Alomone Labs, Jerusalem, Israel), rabbit monoclonal anti-androgen receptor antibody (Cat. No. 5153, Cell Signaling, Danvers, MA), rabbit monoclonal anti-CD11b (Cat. No. 6410, Bio SB, Santa Barbara, CA.), or rabbit monoclonal anti-CD3 (Cat. No. RM-9107-S0, ThermoFisher Scientific, Waltham, MA) for 45 minutes at RT. Poly-HRP–conjugated anti-mouse IgG antibody or anti-rabbit IgG was used as secondary antibody. Staining was visualized using DAB (Cat. No. D4168, Sigma-Aldrich, St. Louis, MO), and slides were counterstained with hematoxylin. Slides were scanned and images processed as described above. For P2X4 image analysis, tumor areas or benign glands were annotated on each TMA spot and the sum intensity per unit area obtained using TMAJ and FrIDA software (version 3.18.0).

IHC and scoring for ERG, PTEN, and TP53 were performed as previously reported in the TMA sets (31,38).

### RISH and IHC Dual stain

RISH staining was performed as described above. After DAB, slides were incubated with mouse monoclonal anti-CD66ce antibody (1 hour), mouse monoclonal anti-tryptase (Cat. No. 2378, Abcam, Cambridge, MA) or mouse monoclonal anti-CD68 (Cat. No. MO814, Dako, Santa Clara, CA) for 45 minutes at RT. Power Vision+ Poly-AP IHC kit was used, followed by Vector red alkaline phosphatase substrate kit (Cat. No. SK-5105, Vector Laboratories, INC. Burlingame, CA) then slides were counterstained with hematoxylin.

### Controls for RISH and IHC

Human-derived prostate cancer PC3 cells which express P2X4 purinergic receptors were incorporated with P2X4 targeted short hairpin RNA (shRNA) plasmids obtained from the JHU ChemCORE (39). Similarly, mouse specific P2X4 shRNAs were incorporated into mouse-derived prostate cancer Myc-CaP cells by lentiviral transduction and puromycin (1ug/mL) selection for at least 5 days. Total protein extracts were obtained and cells were collected into FFPE plugs as previously described (33). These cells were used to confirm specificity of the P2X4 antibodies and RISH probes (Suppl. Fig. S1). FFPE Tonsil tissues were used as positive controls for anti-CD66ce and anti-CD68 IHC (data not shown). Tryptase IHC was previously described (40).

### Mice

8-10 week old male FVB/NJ wildtype mice (Jackson Labs, Bar Harbor, ME) were housed in a specific pathogen-free environment with 12 hour light/dark cycle, and received food and water ad libitum. All procedures were performed under the guidelines of Johns Hopkins Animal Care and Use Committee (ACUC) after a two-week acclimatization period. Animals were sacrificed by CO_2_ asphyxiation followed by exsanguination, and reference tissues along with serum and allograft tumors were collected.

### Allografts

Experimental and control groups were randomly assigned by cage (5 mice per cage, 2 cages per experimental group) and the investigator performing tumor size measurements was blinded to treatment. 1 × 10^6^ Myc-CaP cells in 50% matrigel diluted in PBS were loaded into a 1ml syringe with attached 25 gauge ½ inch needle with care to eliminate air bubbles. 10 week old males were anaesthetized with 2-4% isoflurane and injected subcutaneously with tumor cells into the right side flank. Tumor sizes were measured at 2-3 day intervals using electronic calipers and tumor volumes were calculated as length×width^2^×0.5. Humane endpoints were reached when tumors measured 2.0cm in any direction.

### Statistical Analysis

Data were compared among groups by one-way ANOVA or between groups by two-tailed Mann Whitney U or Wilcoxon matched-pairs signed rank test (Fig. 2-5, Suppl. Fig. 3, 6, & 8).

Spearman’s rank-order correlation was used to compare trends of expression between groups (Fig. 2, 3, Suppl. Fig. 3) and Log-rank test was used to compare survival curves (Fig. 6). Thise data were analyzed and graphed using GraphPad Prism Software (version 7.01; GraphPad Software, Inc., San Diego, CA) and values were considered significant at p < 0.05.

We pooled men from the “Race Disparity” and “PCBN High Grade Race” TMA sets and estimated the median P2X4 protein intensity per unit area of tumor and benign tissue using negative binomial regression and adjusting for TMA set, race/ethnicity, age, stage, grade, PTEN and ERG status (Table 1-2, Suppl. Table S3). Analyses were stratified by PTEN (loss/intact), ERG (negative/positive), grade (≤ 3+4/≥4+3), and race/ethnicity (Black/White). For each tissue type, we categorized men as ≤ the median value of P2X4 (reference) or > the median value, and jointly with PTEN status as ≤ the median value of P2X4 with PTEN intact (reference), ≤ the median value of P2X4 with PTEN loss, > the median value of P2X4 with PTEN intact, or > the median value of P2X4 with PTEN loss. Men on the “Race Disparity” TMA and a subset of men with the appropriate outcomes data on the “PCBN High Grade Race” TMA were pooled to evaluate the association between P2X4 and biochemical recurrence and metastases. Men were followed for a mean of 9.20 and 1.95 years on the “Race Disparity” and “PCBN High Grade Race” TMAs, respectively. We used Cox proportional hazards regression to estimate the relative hazard (HR) and 95% confidence interval (CI) of biochemical recurrence and metastases for P2X4, and for joint biomarker categories of P2X4 and PTEN. Models were adjusted for TMA set, race/ethnicity, age, stage and grade. In sub-analyses, we also adjusted for PTEN. Analyses were conducted using SAS version 9.4.

**Table 1.** Adjusted median P2X4 protein intensity per unit area measurement stratified by PTEN, ERG, and race.*

| **Tissue Type** | **Cancer** |  |  | **Benign** |  |  |
| --- | --- | --- | --- | --- | --- | --- |
| **Gleason** | <=3+4 | >=4+3 | p** | <=3+4 | >=4+3 | p** |
| # cases | 178 | 213 |  | 178 | 213 |  |
| Median | 2.4579 | 2.4927 | 0.9016 | 0.4363 | 0.6965 | 0.0006 |
| **Race** | Black | White | p** | Black | White | p** |
| # cases | 182 | 209 |  | 182 | 209 |  |
| Median | 2.77 | 2.20 | 0.036 | 0.53 | 0.58 | 0.46 |
| **PTEN** | Intact | Loss | p** | Intact | Loss | p** |
| # cases | 302 | 87 |  | 302 | 87 |  |
| Median | 2.16 | 3.33 | 0.0003 | 0.51 | 0.67 | 0.06 |
| **ERG** | Negative | Positive | p** | Negative | Positive | p** |
| # cases | 255 | 134 |  | 255 | 134 |  |
| Median | 1.94 | 3.34 | <.0001 | 0.52 | 0.6 | 0.26 |
| **White Patients** |  |  |  |  |  |  |
| **PTEN** | Intact | Loss | p*** | Intact | Loss | p*** |
| # cases | 134 | 48 |  | 134 | 48 |  |
| Median | 3.13 | 4.32 | 0.037 | 0.84 | 0.94 | 0.52 |
| **ERG** | Negative | Positive | p*** | Negative | Positive | p*** |
| # cases | 95 | 87 |  | 95 | 87 |  |
| Median | 2.3899 | 4.4341 | <.0001 | 0.8577 | 0.8814 | 0.8671 |
| **Black Patients** |  |  |  |  |  |  |
| **PTEN** | Intact | Loss | p*** | Intact | Loss | p*** |
| # cases | 168 | 39 |  | 168 | 39 |  |
| Median | 1.94 | 3.20 | 0.0081 | 0.50 | 0.68 | 0.18 |
| **ERG** | Negative | Positive | p*** | Negative | Positive | p*** |
| # cases | 160 | 47 |  | 160 | 47 |  |
| Median | 1.9704 | 2.9725 | 0.0241 | 0.5101 | 0.6211 | 0.3701 |
| **PTEN Adjusted** |  |  |  |  |  |  |
| **Race** | White | Black | p# | White | Black | p# |
| # cases | 182 | 207 |  | 182 | 207 |  |
| Median | 2.96 | 2.43 | 0.071 | 0.55 | 0.62 | 0.37 |
| **ERG Adjusted** |  |  |  |  |  |  |
| **Race** | White | Black | p## | White | Black | p## |
| # cases | 182 | 207 |  | 182 | 207 |  |
| Median | 2.62 | 2.47 | 0.59 | 0.53 | 0.60 | 0.31 |

\*, participants from “Race Disparity” and “PCBN High Grade Race” TMA sets.
From negative binomial regression with the adjustment for \*\* TMA set, age, race, grade, and stage;
\*\*\*adjustment for TMA set, age, grade, and stage; #TMA set, age, grade, stage, and PTEN status;

**Table 2.** Association of P2X4 expression and joint categories of P2X4 expression and PTEN loss with prostate cancer metastasis and biochemical recurrence.*

|  | Cases (N) | Person-Years | HR (95% CI) ** |  |  | p |
| --- | --- | --- | --- | --- | --- | --- |
| **Metastasis** | *Benign* |  |  |  |  |  |
| Below median | 10 | 861 |  | Ref. |  |  |
| Above median | 25 | 1043 | 2.57 | 1.18 | 5.60 | 0.02 |
| Below median*** | 10 | 861 |  | Ref. |  |  |
| Above median*** | 25 | 1043 | 2.67 | 1.21 | 5.90 | 0.02 |
| Below median /PTEN intact | 7 | 712 |  | Ref. |  |  |
| Below median /PTEN loss | 3 | 149 | 1.01 | 0.23 | 4.36 | 0.99 |
| Above median /PTEN intact | 18 | 769 | 2.39 | 0.92 | 6.23 | 0.08 |
| Above median /PTEN loss | 7 | 274 | 3.36 | 1.01 | 11.2 | 0.05 |
|  | *Cancer* |  |  |  |  |  |
| Below median | 12 | 933 |  | Ref. |  |  |
| Above median | 23 | 971 | 1.88 | 0.88 | 4.00 | 0.10 |
| Below median*** | 12 | 933 |  | Ref. |  |  |
| Above median*** | 23 | 971 | 1.88 | 0.88 | 4.00 | 0.10 |
| Below median /PTEN intact | 11 | 831 |  | Ref. |  |  |
| Below median /PTEN loss | 1 | 101 | 0.25 | 0.03 | 2.23 | 0.22 |
| Above median /PTEN intact | 14 | 649 | 1.25 | 0.51 | 3.05 | 0.62 |
| Above median /PTEN loss | 9 | 322 | 1.83 | 0.67 | 4.97 | 0.24 |
| **Biochemical Recurrence** | *Benign* |  |  |  |  |  |
| Below median | 40 | 702 |  | Ref. |  |  |
| Above median | 61 | 767 | 1.44 | 0.96 | 2.16 | 0.08 |
| Below median*** | 40 | 702 |  | Ref. |  |  |
| Above median*** | 61 | 767 | 1.45 | 0.96 | 2.17 | 0.08 |
| Below median /PTEN intact | 26 | 607 |  | Ref. |  |  |
| Below median /PTEN loss | 14 | 95 | 1.59 | 0.81 | 3.14 | 0.18 |
| Above median /PTEN intact | 39 | 567 | 1.44 | 0.87 | 2.39 | 0.15 |
| Above median /PTEN loss | 22 | 200 | 2.31 | 1.26 | 4.24 | 0.01 |
|  | *Cancer* |  |  |  |  |  |
| Below median | 47 | 729 |  | Ref. |  |  |
| Above median | 54 | 740 | 1.27 | 0.84 | 1.91 | 0.25 |
| Below median*** | 47 | 729 |  | Ref. |  |  |
| Above median*** | 54 | 740 | 1.19 | 0.79 | 1.8 | 0.41 |
| Below median /PTEN intact | 34 | 668 |  | Ref. |  |  |
| Below median /PTEN loss | 13 | 61 | 1.84 | 0.90 | 3.8 | 0.09 |
| Above median /PTEN intact | 31 | 506 | 1.311 | 0.79 | 2.2 | 0.30 |
| Above median /PTEN loss | 23 | 234 | 1.82 | 1.03 | 3.21 | 0.04 |

\*, participants from Race Disparity” and “PCBN High Grade Race” TMA sets. Median was determined based on the P2X4 measurement among all men. Hazard ratio (HR) and 95% confidence intervals (CI) estimated from Cox Hazard regression model adjusting for \*\*age, stage, grade, race, and TMA sets;
\*\*\*age, stage, grade, race, TMA sets, and PTEN status.

## Results

### P2X4 purinergic receptor is overexpressed in prostate cancer

We began by examining all 15 human P2 purinergic receptors in an RNAsequencing dataset (n=25) obtained on PCa and matched benign tissues (41). We identified P2X4 purinergic receptor expression as elevated in cancer compared to benign tissues (p = 0.0045) in both low grade (p = 0.0168) and high grade (Gleason sum ≥ 4 +3) cases (p = 0.0474, Suppl. Fig. S2A). P2X4 overexpression in PCa was corroborated in publicly available gene datasets, Tomlins (p = 7.44E-5), Lapointe (p = 8.81E-5), Vanaja (p = 4.07E-5), Welsh (p = 1.93E-5), and Wallace (p = 0.002) analyzed via ONCOMINE (Suppl. Fig. S2B-F) (42-47). P2X4 receptor mRNA expression was also elevated in PIN compared to benign tissues (p = 9.48E-4) in the Tomlins dataset (Suppl. Fig. S2B) (42). Analysis of TCGA RNA sequencing data via Wanderer (48) further corroborated elevated P2X4 receptor expression in PCa (n = 374) compared to benign tissues (n = 52, p = 9.10E-12, Suppl. Fig. S2G).

### P2X4 purinergic receptor protein is primarily expressed in epithelial cells and immune cells in the prostate

We next aimed to determine the cellular localization and distribution of P2X4 receptor expression in normal prostate tissues. IHC analysis was performed on FFPE whole tissue sections from normal organ donor prostate tissues with no cancer (n=4, Suppl. Table S1). PX4 receptor expression was observed primarily in epithelial cells of the prostate gland. Prostate-infiltrating immune cells had more intense P2X4 expression than epithelium, particularly in regions of inflammation (Fig. 1A). Next, we performed P2X4 IHC on FFPE whole tissue sections from a representative tumor-containing block, a matched benign tissue block with inflammation, and a matched benign tissue block without overt inflammation in radical prostatectomy specimens from 3 Black and 3 White patients (Suppl. Table S1). Similar to organ donor prostates, there was P2X4 receptor protein expression in epithelial cells of benign glands (Fig. 1B, C). We observed increased intensity of P2X4 receptor expression in regions of PIN and cancer compared to benign glands (Fig. 1B, C). There was also intense P2X4 receptor expression observed on immune cells in the stromal compartment and in the lumens of glands (Fig. 1D). Dual IHC staining indicated that tryptase-positive mast cells do not express P2X4 receptors (Fig. 1E). On the contrary, some CD66ce^+^ myeloid cells, considered in most cases to be neutrophils, and most CD68^+^ macrophages expressed P2X4 receptors (Fig. 1E).

**Figure 1:**
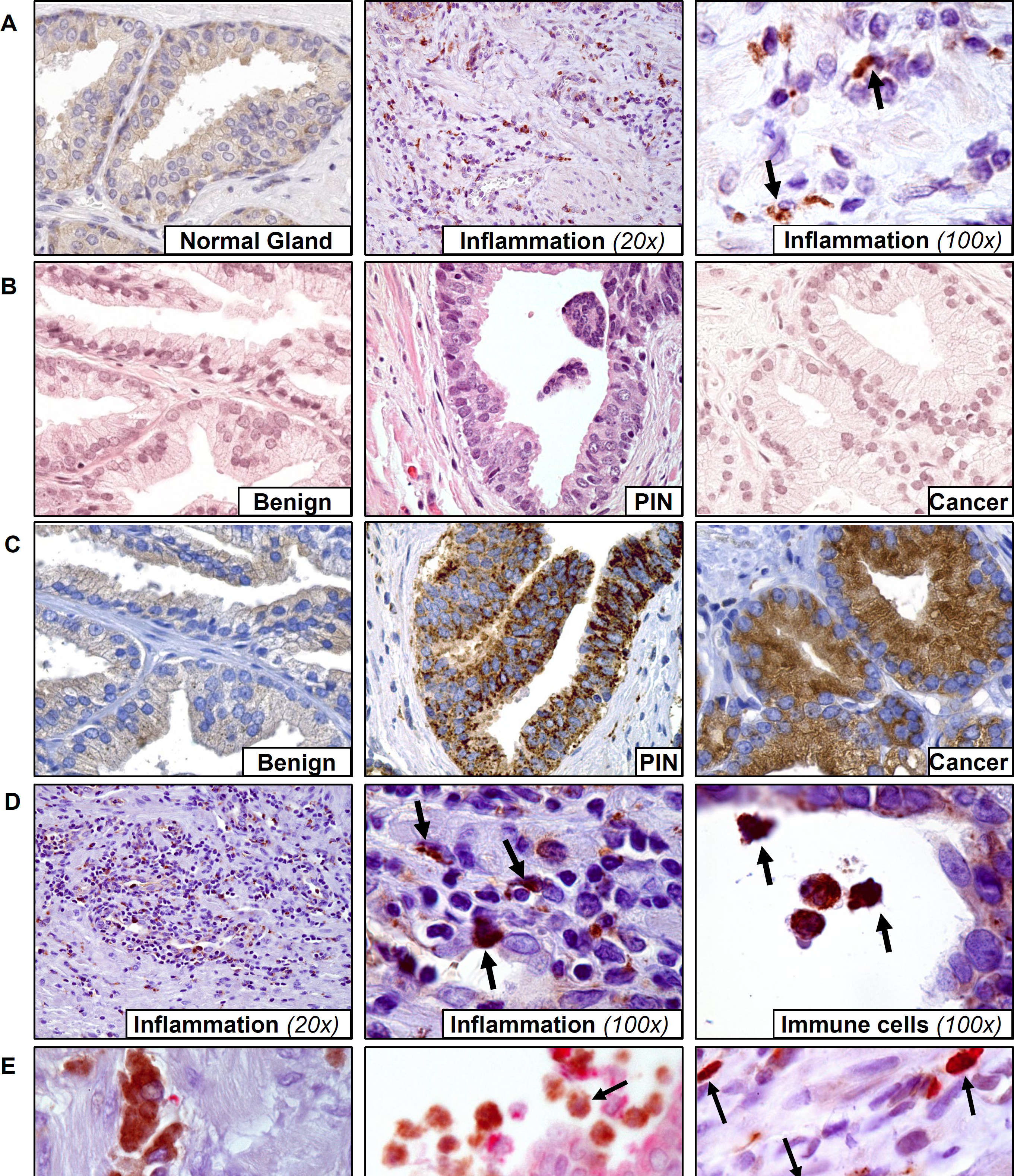
P2X4 purinergic receptor protein is expressed (A) modestly in the epithelium of prostate glands and more intensely on immune cells of organ donor prostates (C) modestly in benign glands and more intensely in prostatic intraepithelial neoplasia (PIN) and cancer in prostatectomy samples and (D) on immune cells. (B) H&E staining confirmed tissue types. (E) Dual staining showed that tryptase positive mast cells (red) do not express P2X4 protein (brown), but some CD66 positive neutrophils (brown) and most CD68 positive macrophages (brown) express P2X4 receptors (red). Images are at 40x magnification unless otherwise stated.

**Figure 2.**
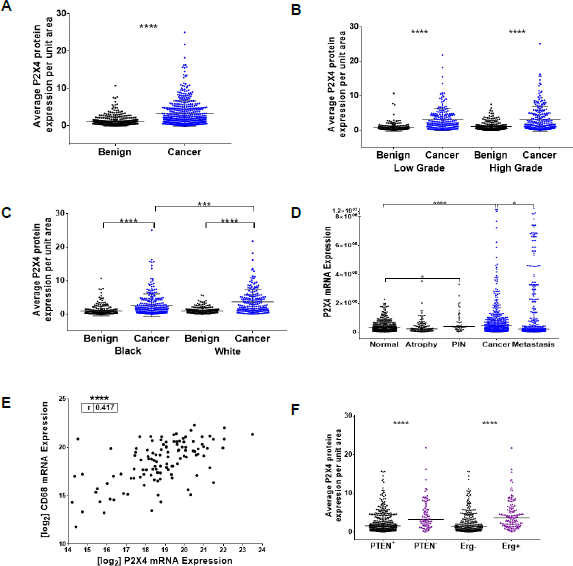
There is elevated P2X4 receptor protein expression in (A) cancer compared to benign tissues (n = 456; p < 0.0001) (B) from both high grade and low grade cases, and (C) in cancer spots from European American men compared to cancer spots from African American men. There is elevated P2X4 mRNA expression in prostatic intraepithelial neoplasia (PIN) and cancer while there is a wide range of expression across metastatic sites. (E) There was elevated P2X4 protein expression in tumors with Erg expression and in tumors with PTEN loss. (F) There is a positive correlation between P2X4 mRNA expression and CD68 mRNA expression.

**Figure 3.**
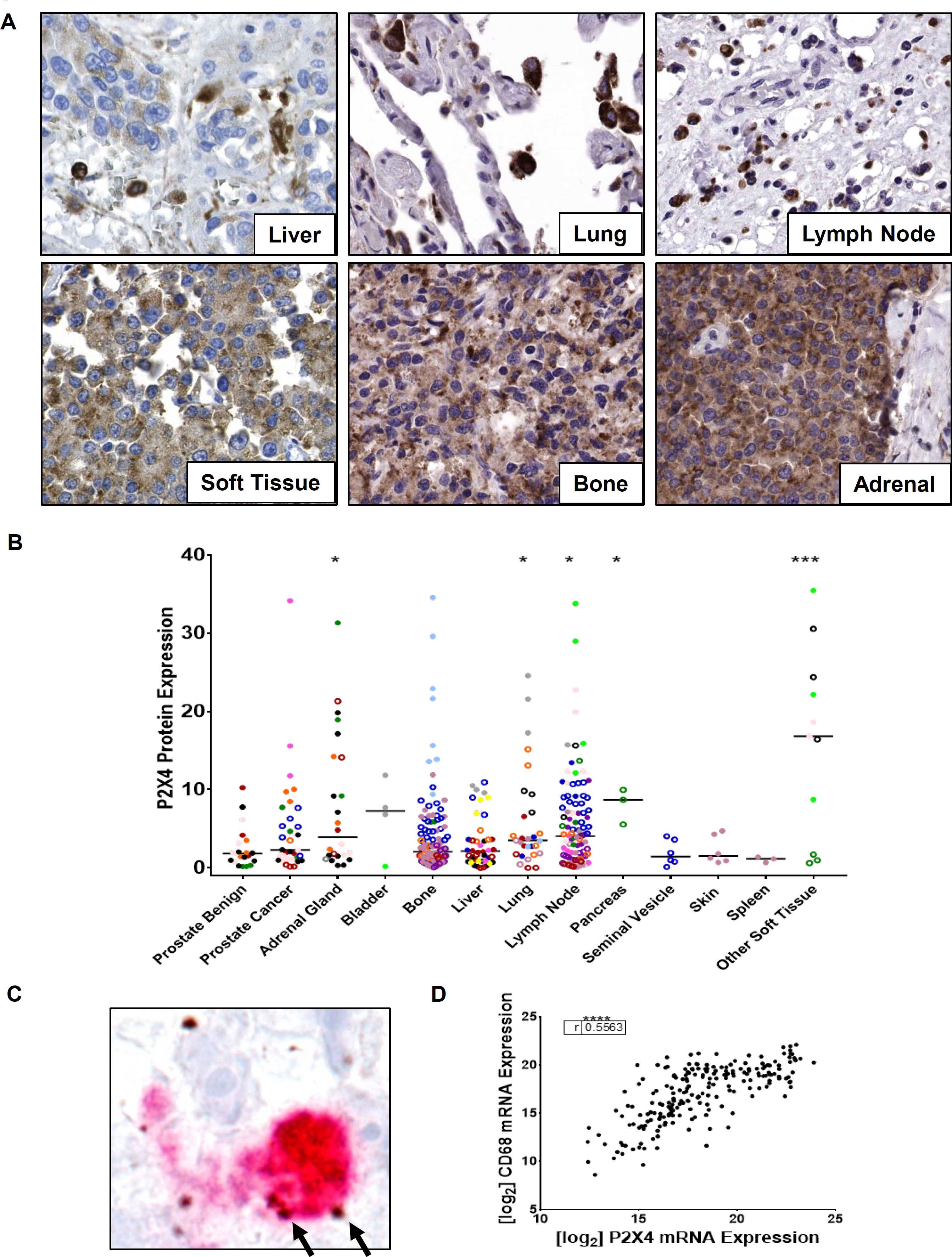
(A, B) P2X4 mRNA expression was detected in cancer and immune cells and in various metastatic sites. (C) P2X4 mRNA expression (brown) detected on CD68^+^ (red) macrophages in metastatic tissues. (D) There is a positive correlation between CD68 and P2X4 mRNA expression in metastatic tissues.

**Figure 4.**
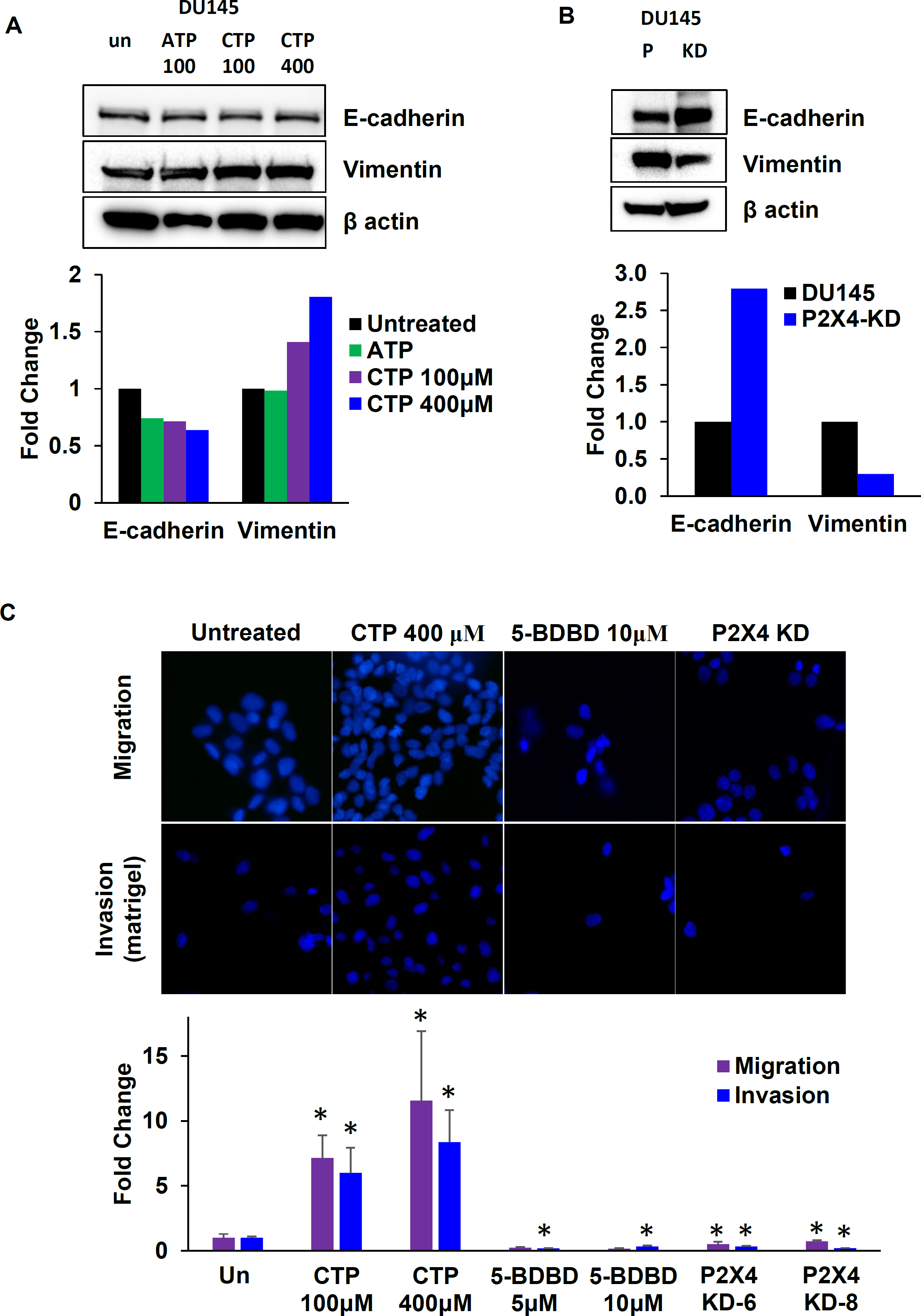
(A) Nucleotide treatment and (B) P2X4 knock down in DU145 cells. (C) CTP treatment increased cell migration and invasion while 5-BDBD and P2X4 knock down decreased cell migration and invasion in transwell assays. (P = parental, KD = knockdown)

**Figure 5.**
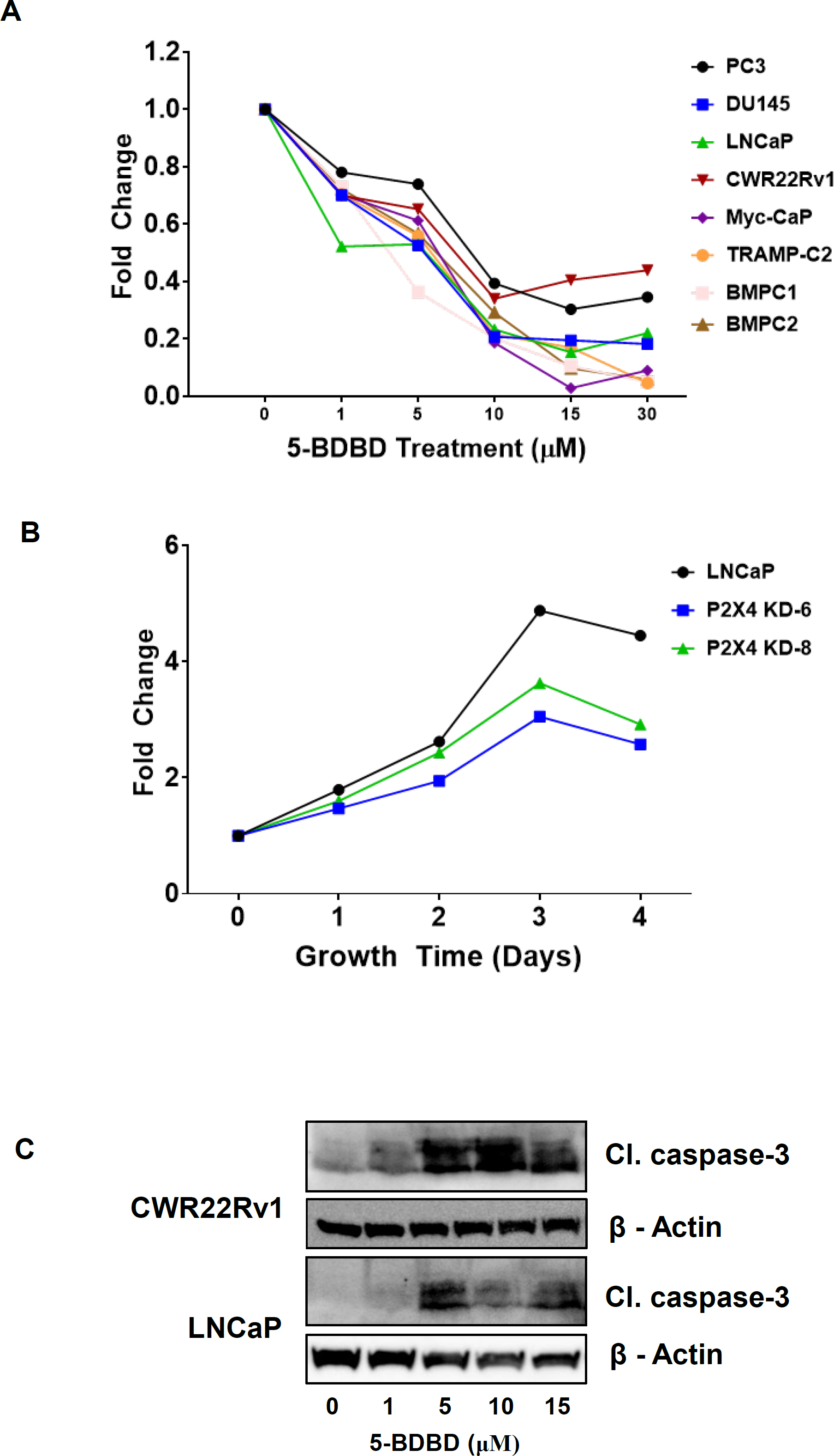
(A) P2X4 antagonist 5-BDBD resulted in a dose-dependent decrease in PCa cell viability. (B) P2X4 down in LNCaP cells resulted attenuated cell growth over time. (C) Treatment with 5-BDBD resulted in a dose-dependent induction of cleaved caspase-3 in PCa cells.

**Figure 6.**
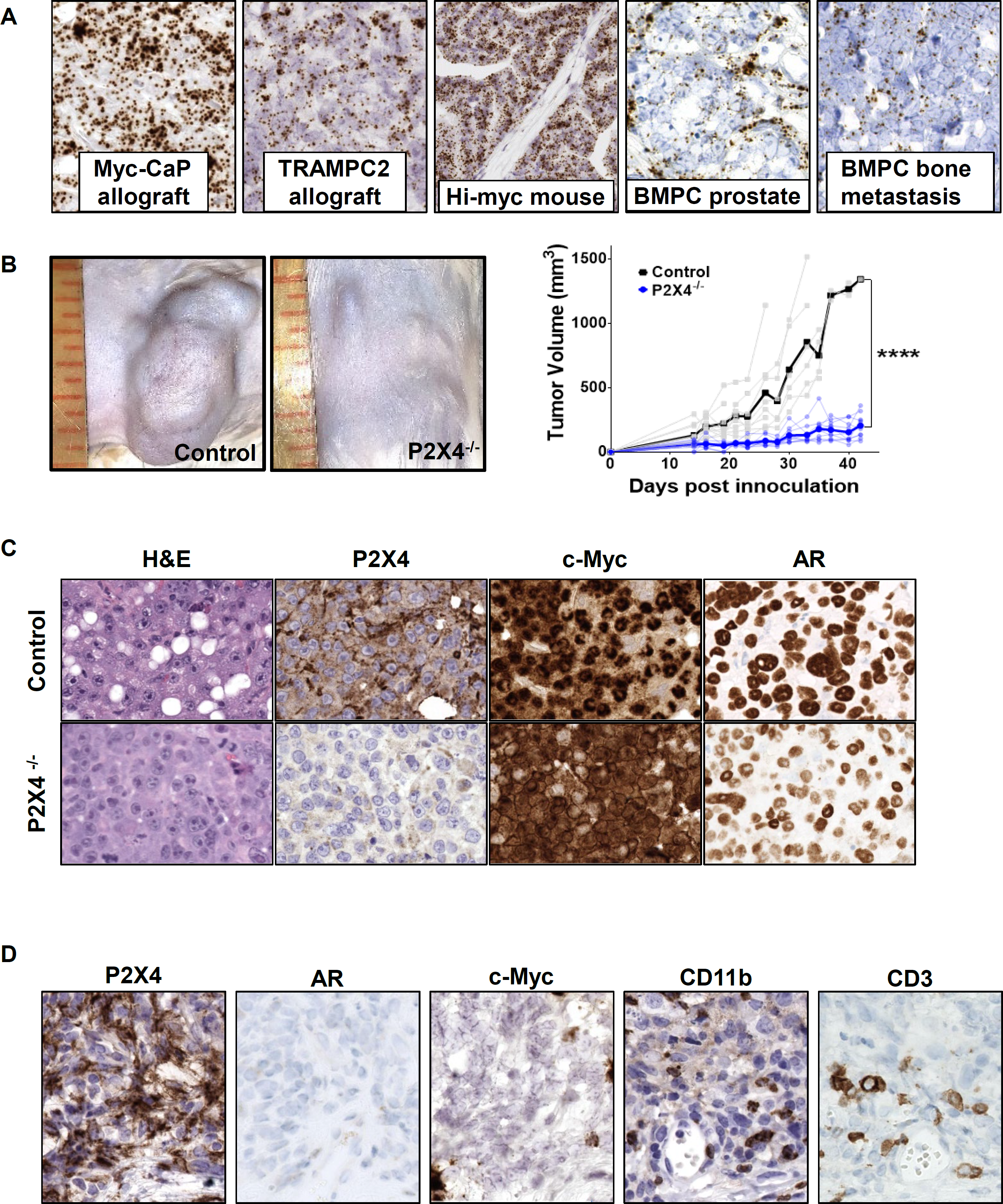
(A) Robust P2X4 mRNA expression is detected in mouse prostate tumors. (B) CRISPR knockdown of P2X4 receptor expression in Myc-CaP cells (n=10) resulted in attenuated allograft growth in FVB/NJ mice compared to control Myc-CaP cells (n=10). (C) Human c-Myc RISH and P2X4 and AR IHC analysis confirmed P2X4 reduced expression in Myc-CaP allograft tissue from mice. (D) P2X4 positive regions lacking AR and human c-Myc stain positive for CD11b and CD3 in Myc-CaP allograft tissue from mice.

### P2X4 purinergic receptor expression is elevated in PIN, prostate cancer, and inflamed regions with increased CD68^+^ macrophages

We next quantified P2X4 receptor protein expression in PCa TMA sets (n = 491 cases, Suppl. Table S2). There was significantly elevated P2X4 receptor expression in cancer regions compared to benign regions (Wilcoxon matched-pairs signed rank analysis, p < 0.0001, Fig. 2A), in both low grade (p < 0.0001) and high grade (p < 0.0001) tissues (Fig. 2B). P2X4 protein expression was significantly increased in benign tissues from higher grade compared to low grade cases (Table 1). P2X4 expression was increased in cancer versus benign tissues from both Black men (p < 0.0001) and White men (p < 0.0001, Fig. 2C). Interestingly, there was significantly higher P2X4 receptor protein expression in cancer tissues from White men compared to cancer tissues from Black men (p = 0.0006, Fig. 2C, Table 1).

RISH analysis of *P2X4* receptor mRNA in the PCBN High Grade Race TMA set (n=120) also showed elevated expression in cancer compared to normal appearing benign tissues (p < 0.0001, Fig 2D). Benign tissue spots on the TMA were further classified as normal (normal appearing, non-inflamed) or as containing atrophy or PIN. There was increased P2X4 receptor mRNA expression in PIN compared to normal benign tissues (p = 0.0152, Fig 2D), corroborating the Tomlins data (Suppl. Fig. S2) (42). There was a variable degree of P2X4 receptor mRNA expression in metastatic tissues collected at autopsy but overall there was significantly more P2X4 in cancer tissues compared to metastatic tissues (p < 0.05, Fig. 2D, Suppl. Table S2).

Based on our observation that CD68^+^ macrophages and some CD66ce^+^ neutrophils express P2X4 receptors, we quantified CD68 mRNA expression and CD66ce protein expression in the PCBN High Grade Race TMA set. Interestingly, we found a positive correlation between P2X4 mRNA and CD68 mRNA expression (r = 0.4084, p < 0.0001, Fig, 2E) but not between P2X4 mRNA and CD66ce protein expression (Suppl. Fig. S3A).

### P2X4 receptor protein expression is elevated in prostate cancer cases with ERG positivity or PTEN loss

We stratified cancer cases based on common genetic alterations of *PTEN* loss, *ERG* positivity, or *TP53* missense mutation. *PTEN* is the most commonly inactivated tumor suppressor in PCa and *PTEN* loss is associated with aggressive PCa (49). *ERG* expression, observed in about 50% of PCa, is commonly observed in cases with *PTEN* loss (49). *TP53* alterations occur in about 53.3% of metastatic castration resistant prostate cancer (mCRPC) and is among the most frequently aberrant genes in mCRPC cases (50). Our analysis revealed that cancer tissues with loss of PTEN protein expression had significantly elevated P2X4 receptor expression compared to normal PTEN expressing cancer tissues (p < 0.0001, Fig. 2F, Table 1). Similarly, cancer tissues with ERG protein expression had significantly higher P2X4 receptor expression compared to ERG-negative cancer tissues (p < 0.001, Fig. 2F, Table 1). There was no difference in P2X4 receptor expression between cancer tissues with or without TP53 nuclear accumulation (Suppl. Fig. S3B). PTEN loss and ERG positivity are more frequent in White compared to Black men (30). To do determine whether the dissimilar frequency of PTEN or ERG alterations between races contributed to increased P2X4 expression observed among White men, we assessed P2X4 expression by PTEN and ERG status separately by race. P2X4 expression was significantly elevated in cases with PTEN loss and ERG positivity in both White and Black men (Table 1). Interestingly, statistical adjustment for PTEN or ERG status each attenuated the difference in P2X4 expression observed between races (Table 1).

### Elevated P2X4 purinergic receptor expression is associated with increased risk of metastasis

We performed P2X4 IHC analysis in the Metastasis at Autopsy TMA. P2X4 protein was expressed on immune cells in the metastatic niche and in metastatic cancer cells (Fig. 3A). We quantified a broad range of expression across patients and between sites within the same patient (Fig. 3B). The mean P2X4 receptor protein expression from adrenal gland, lung, lymph node, and pancreas were each significantly elevated compared to prostate benign tissues (harvested at autopsy e.g., the men had not undergone radical prostatectomy) (Fig. 3B). Dual staining revealed positive P2X4 mRNA expression on CD68^+^ macrophages in metastatic tissues (Fig. 3C). Similar to primary PCa tissues, there was a positive correlation between P2X4 mRNA and CD68 mRNA expression in the Metastasis at Autopsy TMA set (r = 0.5563, p < 0001, Fig. 3D).

Compared to men with P2X4 below the median in benign glands, men with P2X4 protein expression above the median had a greater than two-fold increased risk of metastasis (HR = 2.57, 95% CI:1.18, 5.60), and a suggestive increased risk of biochemical recurrence (HR = 1.437, 95% CI: 0.96, 2.16) (Table 2). PTEN loss has been shown to be prognostic in prostate cancer (30,51), however, adjustment for PTEN did not change inferences (Table 2). We examined the combined effect of P2X4 receptor expression in benign tissues and PTEN loss. As compared to men with P2X4 below the median and PTEN intact, men with P2X4 above the median and PTEN loss had a greater than 3-fold increased risk of metastasis (HR = 3.36, 95% CI: 1.01,11.18), and greater than two-fold increased risk of biochemical recurrence (HR = 2.3, 95% CI:1.26, 4.24) (Table 2). Interestingly, as compared to men with P2X4 expression in cancer below the median and PTEN intact in cancer tissue, men with P2X4 expression in cancer above the median and PTEN loss had a significantly increased risk of biochemical recurrence (HR = 1.82, 95% CI: 1.03, 3.21; Table 2). P2X4 expression in cancer tissue was not significantly associated with metastasis (Table 2).

### P2X4 purinergic receptor activation increases prostate cancer cell migration and invasion

RISH and IHC analysis detected P2X4 receptor expression in human derived PCa cell lines PC3, DU145, LNCaP, CWR22Rv1, VCaP, LAPC4, and MD PCa 2b (Suppl. Fig S4). Similarly, P2X4 receptor protein and mRNA expression were detected in mouse derived BMPC1, BMPC2, Myc-CaP, and TRAMP-C2 cell lines (Suppl. Fig. S5). Considering the association between P2X4 receptor expression and increased risk of metastasis, we assessed the effect of P2X4 receptor activation on the migratory or invasive phenotype of PCa cells. Treatment of PC3 cells with endogenous broad spectrum P2 receptor ligand ATP (100μM) or P2X4 selective agonist CTP (100μM) modestly decreased expression of epithelial markers E-cadherin and cytokeratin 8 by western blotting (WB) analysis (Suppl. Fig. S6A). Concurrently, ATP and CTP treatment increased expression of mesenchymal markers vimentin and snail, suggesting that P2X4 activation may promote a migratory phenotype in PCa cells (Suppl. Fig. S6A). Similar results were seen in E-cadherin and vimentin WB analysis of DU145 cells (Fig. 4A). We used CRISPR-cas9 gene editing to knock down P2X4 expression in DU145 cells (Suppl. Fig. S7A). P2X4 knockdown (KD) resulted in increased e-cadherin and decreased vimentin protein expression in DU145 cells (Fig. 4B).We performed functional transwell assays to assess the effect of P2X4 activation on cell migration and invasion. Nucleotide treatment alone was sufficient to significantly increase PC3 and CWR22Rv1 cell migration across transwell membranes and invasion across matrigel coated transwell membranes at 24h, 48h, or 72h (Suppl. Fig S6B).

Specifically, CTP (100, 400μM) 72h treatment significantly increased both migration (7.2-fold, p=0.018; 11.6-fold, p=0.04) and invasion (6.0-fold, p=0.045; 8.4-fold, p=0.009) of DU145 cells (Fig. 4C). Conversely, treatment with P2X4 selective antagonist, 5-BDBD (5, 10μM) significantly decreased DU145 cell migration (0.2-fold, p=0.001; 0.2-fold, p=0.008) and invasion (0.2-fold, p<0.001; 0.3-fold, p<0.0001, Fig. 4B). Further, P2X4 KD also resulted in reduced cell migration (0.5-fold, n.s.) and invasion (0.3-fold, p=0.001) in DU145 cells (Fig. 4C).

### P2X4 purinergic receptor blockade decreases prostate cancer cell viability

We next assessed the effect of P2X4 blockade on PCa cell viability. 5-BDBD treatment resulted in a dose-dependent decrease in cell viability of PC3, DU145, LNCaP, CWR22Rv1, TRAMP-C2, Myc-CaP, BMPC1, and BMPC2 cells as assessed by MTT assay (Fig. 5A, Suppl. Fig. S8A). P2X4 KD attenuated growth rates of LNCaP and DU145 cells (Fig. 5B, Suppl. Fig. S8B). 5-BDBD treatment induced dose dependent expression of apoptotic marker cleaved caspase-3 in LNCaP and CWR22Rv1 cells (Fig. 5C). Temporary knockdown of P2X4 by siRNA also induced cleaved caspase-3 expression in PC3 cells (Suppl. Fig. S8C).

### P2X4 purinergic receptor knockdown attenuates Myc-CaP allograft tumor growth

We next assessed whether P2X4 receptor function affects tumor initiation and progression *in vivo*. We confirmed P2X4 receptor expression in Myc-CaP allograft, TRAMP-C2 allograft, Hi-Myc prostate tumor, BMPC prostate tumor, and BMPC bone metastasis tissues. We observed robust P2X4 mRNA in all four models (Fig. 6A). We selected an allograft model for the feasibility of genetically manipulating P2X4 expression directly in cancer cells and the benefit of using an immunocompetent host. We used CRISPR-cas9 to knock down P2X4 receptor expression in Myc-CaP cells. Myc-CaP control (CRISPR empty vector) and Myc-CaP P2X4 KD cells in 50% matrigel were subcutaneously allografted into 10-week old male FVB/NJ mice (n=10). Cages were blinded to the investigator that measured tumor growth. Myc-CaP-P2X4 KD cells resulted in significantly attenuated subcutaneous allograft growth in FVB/NJ mice compared to control Myc-CaP cells (Fig. 6B). Tumors were palpable or measurable in all 10 mice in the control group, compared to 8 out of 10 in the P2X4 KD group. The tumors that were established in the P2X4 KD group progressed significantly slower than tumors in the control group (p < 0.0001, Fig. 6B). H&E staining, RISH analysis of human c-Myc, and IHC analysis of AR were used to confirm that tumors were developed from allografted Myc-CaP cells (Fig. 6C). IHC analysis of P2X4 confirmed P2X4 reduction in allografts generated from Myc-CaP-P2X4 KD cells (Fig. 6C). Interestingly, we observed substantial P2X4 protein expression in some regions lacking human c-Myc mRNA and AR protein expression (Fig. 6D). These regions likely represented the host tumor microenvironment (TME). IHC analysis identified CD11b^+^ immune cells, likely macrophages or neutrophils, and CD3^+^ T cells in these P2X4 positive non-tumor regions (Fig. 6D, Suppl. Fig. 9).

## Discussion

Herein we report a comprehensive profile of P2X4 receptor expression in normal prostate, primary PCa, metastatic PCa, human and mouse PCa cells, and mouse PCa tissues. We demonstrate P2X4 receptor protein expression in the epithelium and infiltrating immune cells of the prostate and measured elevated P2X4 receptor expression in human PIN and cancer by RNA sequencing analysis, RISH, and IHC. Higher P2X4 protein expression was observed in cases with PTEN protein loss and cases with positive ERG protein expression. Higher P2X4 protein expression was also associated with an increased risk of metastasis and above median P2X4 protein expression coupled with PTEN loss was associated with an increased risk of both metastasis and biochemical recurrence after prostatectomy.

P2X4 purinergic receptors have primarily been studied for their role in neuropathic and inflammatory pain (52,53). In fact, a P2X4 antagonist developed for the treatment of neuropathic pain has completed a Phase 1 clinical trial (54,55). Like other P2X purinergic receptors, P2X4 forms heterotrimers with P2X2, P2X5, and/or P2X6 and is localized at the plasma membrane (56). P2X4 is unique in that it may also localize to intracellular compartments such as lysosomes (56). Our IHC analysis demonstrates that P2X4 receptor expression is not limited to the plasma membrane in PCa. While P2 purinergic receptors have been widely investigated in different cancer sites, studies on P2X4 receptor expression and function in cancer have rarely been reported (20,21,24-28). A recent study identified the P2X4 purinergic receptor as a potential clinical target for patients with PCa (29). The authors demonstrate that pharmacological inhibition of P2X4 receptors impaired PCa cell growth and decreased PCa cell motility (29).

Our study corroborates these findings using additional cell lines and using genetic manipulation to confirm the specific role of P2X4 purinergic receptors in PCa cell viability, migration and invasion. Further, we report that direct activation of P2X4 receptors alone was sufficient to promote PCa migration and invasion in transwell assays. Future studies will be aimed at determining the mechanistic pathways through which P2X4 receptors function in PCa cells.

Our dual staining analysis identified some CD66^+^ neutrophils, most CD68^+^ macrophages, but no tryptase^+^ mast cells as P2X4 receptor-expressing immune cells in PCa tissues. We also found a positive correlation between P2X4 mRNA and CD68 mRNA expression in the PCBN High Grade and Metastasis at Autopsy TMA sets. Considering the high amounts of P2X4 receptor expression we observe and measure in epithelial cells, we believe that this positive correlation is not primarily due to macrophages expressing P2X4 receptors. Rather, there may be higher macrophage infiltration in regions with high P2X4 receptor expression. A study reports that 5-BDBD treatment in a mouse model of ischemic stroke reduced infiltration of total leukocytes including neutrophils and monocytes into the brain after stroke (57). Other studies report P2X4 receptors as involved in macrophage activation, facilitating macrophage phagocytosis and

NLRP1/3 inflammasome induction (56,58,59). High numbers of M2 macrophages are associated with poorer outcomes in men with PCa (60,61). As such, investigating P2X4 receptor function in the context of PCa would be worthwhile.

We report higher P2X4 receptor expression in cases with PTEN loss and cases with high P2X4 expression and PTEN loss have an increased risk of metastasis and biochemical recurrence.

Future studies will be aimed at assessing the feasibility of using P2X4 receptor expression in combination with other markers as a prognostic tool. Additionally, the biological associations of P2X4 receptors and PTEN is not yet fully investigated. PTEN is a negative regulator of PI3K resulting in de-phosphorylation of Akt (62). Phosphorylated Akt (p-Akt), its active form, is associated with poor prognosis in multiple cancers and promotes mesenchymal-like properties and metastasis of PCa cells (63). Interestingly, a study reported crosstalk between PTEN and another Akt regulatory phosphatase, PHLPP that is dependent on P2X4 receptor signaling (62). The authors propose that this crosstalk controls the levels of p-Akt, promotes PCa cell invasion, and is mediated by the P2X4 purinergic receptor. Further, this study showed that the P2X4 receptor is necessary for TGF-β mediated PC3 cell invasiveness (62).

Higher P2X4 receptor expression was also measure in cases with ERG positivity within our cohorts. ERG positivity in PCa is typically the result of gene fusions between the androgen-regulated gene *TMPRSS2* and the transcription factor *ERG* (64). One study reported the P2Y2 purinergic receptor among common upregulated genes in TMPRSS2:ERG-expressing PC3 cells (65). The study found that the collective TMPRSS2:ERG target genes were associated with PC3 cell motility and invasiveness, further corroborating the role for the P2 purinergic receptor family in driving PCa cell invasiveness (65).

Our study demonstrated that P2X4 receptors are involved in the tumor initiation and progression of PCa subcutaneous allografts in mice. These data provide a strong proof of concept that P2X4 purinergic receptors, specifically, have a direct role in tumor development. The Myc-CaP allograft model allowed us to assess tumor growth in a fully immunocompetent mouse. Our histological analysis demonstrated that P2X4 receptor expression was robust on immune cells and throughout the TME. Future studies will investigate the role of P2X4 purinergic receptors in the PCa TME.

Overall, our study demonstrates a role for P2X4 purinergic receptors in PCa aggressiveness. The associations of P2X4 receptor expression with common genetic alterations and increased risk of metastasis or biochemical recurrence highlight the P2X4 purinergic receptor as a potential prognostic factor. Further, the current study indicates a functional role for P2X4 in maintaining PCa cell viability, promoting PCa cell motility and invasiveness, and driving PCa tumor development *in vivo*. These data and the existence of a P2X4 antagonist in-development highlight the P2X4 purinergic receptor a therapeutic target for the treatment of aggressive PCa.

## ADDITIONAL INFORMATION

### Authors’ contributions

JPM and KSS conceived experiments. IV, LBP, TLL, and AMD are pathologists that characterized all patient and mouse tissues and assisted with construction of TMAs. JPM, JH, LM, TA, RK, and AMC carried out experiments and JPM, TA, RS, and KSS analyzed the data. JPM, JL, and CEJ performed statistical analyses. JPM and KSS were involved in writing the manuscript and all authors assisted with editing the manuscript and had final approval of the submitted and published versions.

## Acknowledgements

The authors thank the patients and their families who participated in the studies at Johns Hopkins. We also thank students Julia Flores and Xavier A. Avilés for their contributions to the lab.

## References

1. Howlader N. NAM, Krapcho M., Miller D., Bishop K., Altekruse S.F., Kosary C.L., Yu M., Ruhl J., Tatalovich Z., Mariotto A., Lewis D.R., Chen H.S., Feuer E.J., Cronin K.A. (eds). SEER Cancer Statistics Review, 1975–2013. National Cancer Institute Bethesda, MD 2015

2. Odedina FT, Akinremi TO, Chinegwundoh F, Roberts R, Yu D, Reams RR, et al. Prostate cancer disparities in Black men of African descent: a comparative literature review of prostate cancer burden among Black men in the United States, Caribbean, United Kingdom, and West Africa. Infectious Agents and Cancer 2009;4:S2

3. Sfanos KS, Yegnasubramanian S, Nelson WG, De Marzo AM. The inflammatory microenvironment and microbiome in prostate cancer development. Nature Reviews Urology 2018;15:11–24

4. De Marzo AM, Platz EA, Sutcliffe S, Xu J, Gronberg H, Drake CG, et al. Inflammation in prostate carcinogenesis. Nat Rev Cancer 2007;7:256–69

5. Sfanos KS, Yegnasubramanian S, Nelson WG, De Marzo AM. The inflammatory microenvironment and microbiome in prostate cancer development. Nat Rev Urol 2018;15:11–24

6. Elliott MR, Chekeni FB, Trampont PC, Lazarowski ER, Kadl A, Walk SF, et al. Nucleotides released by apoptotic cells act as a find-me signal to promote phagocytic clearance. Nature 2009;461:282

7. Hernandez C, Huebener P, Schwabe RF. Damage-associated molecular patterns in cancer: a double-edged sword. Oncogene 2016;35:5931

8. Chen Y, Corriden R, Inoue Y, Yip L, Hashiguchi N, Zinkernagel A, et al. ATP release guides neutrophil chemotaxis via P2Y2 and A3 receptors. Science (New York, NY) 2006;314:1792–5

9. Kukulski F, Ben Yebdri F, Lecka J, Kauffenstein G, Lévesque SA, Martín-Satué M, et al. Extracellular ATP and P2 receptors are required for IL-8 to induce neutrophil migration. Cytokine 2009;46:166–70

10. Lim HM, Woon H, Han JW, Baba Y, Kurosaki T, Lee MG, et al. UDP-Induced Phagocytosis and ATP-Stimulated Chemotactic Migration Are Impaired in STIM1(-/-) Microglia In Vitro and In Vivo. Mediators of inflammation 2017;2017:8158514

11. Burnstock G. Purinergic signalling: Its unpopular beginning, its acceptance and its exciting future. BioEssays 2012;34:218–25

12. von Kügelgen I, Wetter A. Molecular pharmacology of P2Y-receptors. Naunyn-Schmiedeberg’s Archives of Pharmacology 2000;362:310–23

13. Idzko M, Ferrari D, Eltzschig HK. Nucleotide signalling during inflammation. Nature 2014;509:310–7

14. Gallucci S, Matzinger P. Danger signals: SOS to the immune system. Current Opinion in Immunology 2001;13:114–9

15. Bours MJL, Swennen ELR, Di Virgilio F, Cronstein BN, Dagnelie PC. Adenosine 5′-triphosphate and adenosine as endogenous signaling molecules in immunity and inflammation. Pharmacology & Therapeutics 2006;112:358–404

16. Burnstock G. P2X ion channel receptors and inflammation. Purinergic Signalling 2016;12:59–67

17. Burnstock G, Di Virgilio F. Purinergic signalling and cancer. Purinergic Signalling 2013;9:491–540

18. White N, Burnstock G. P2 receptors and cancer. Trends in Pharmacological Sciences 2006;27:211–7

19. Burnstock G, Verkhratsky A. Long-term (trophic) purinergic signalling: purinoceptors control cell proliferation, differentiation and death. Cell Death & Disease 2010;1:e9

20. Li W-H, Qiu Y, Zhang H-Q, Tian X-X, Fang W-G. P2Y2 Receptor and EGFR Cooperate to Promote Prostate Cancer Cell Invasion via ERK1/2 Pathway. PLoS ONE 2015;10:e0133165

21. Qiu Y, Li W-h, Zhang H-q, Liu Y, Tian X-X, Fang W-G. P2X7 Mediates ATP-Driven Invasiveness in Prostate Cancer Cells. PLoS ONE 2014;9:e114371

22. Pellegatti P, Raffaghello L, Bianchi G, Piccardi F, Pistoia V, Di Virgilio F. Increased Level of Extracellular ATP at Tumor Sites: In Vivo Imaging with Plasma Membrane Luciferase. PLoS ONE 2008;3:e2599

23. Falzoni S, Donvito G, Virgilio FD. Detecting adenosine triphosphate in the pericellular space. Interface Focus 2013;3:20120101

24. Pellegatti P, Raffaghello L, Bianchi G, Piccardi F, Pistoia V, Di Virgilio F. Increased Level of Extracellular ATP at Tumor Sites: <italic>In Vivo</italic> Imaging with Plasma Membrane Luciferase. PLoS ONE 2008;3:e2599

25. Nylund G, Hultman L, Nordgren S, Delbro DS. P2Y2- and P2Y4 purinergic receptors are over-expressed in human colon cancer. Autonomic and Autacoid Pharmacology 2007;27:79–84

26. Künzli BM, Berberat PO, Giese T, Csizmadia E, Kaczmarek E, Baker C, et al. Upregulation of CD39/NTPDases and P2 receptors in human pancreatic disease. American Journal of Physiology - Gastrointestinal and Liver Physiology 2007;292:G223–G30

27. Maynard JP, Lee J-S, Sohn BH, Yu X, Lopez-Terrada D, Finegold MJ, et al. P2X3 purinergic receptor overexpression is associated with poor recurrence-free survival in hepatocellular carcinoma patients. 2015.

28. Liu Z, Liu Y, Xu L, An H, Chang Y, Yang Y, et al. P2X7 receptor predicts postoperative cancer-specific survival of patients with clear-cell renal cell carcinoma. Cancer science 2015;106:1224–31

29. He J, Zhou Y, Arredondo Carrera HM, Sprules A, Neagu R, Zarkesh SA, et al. Inhibiting the P2X4 Receptor Suppresses Prostate Cancer Growth In Vitro and In Vivo, Suggesting a Potential Clinical Target. Cells 2020;9

30. Tosoian JJ, Almutairi F, Morais CL, Glavaris S, Hicks J, Sundi D, et al. Prevalence and Prognostic Significance of PTEN Loss in African-American and European-American Men Undergoing Radical Prostatectomy. Eur Urol 2017;71:697–700

31. Kaur HB, Guedes LB, Lu J, Maldonado L, Reitz L, Barber JR, et al. Association of tumor-infiltrating T-cell density with molecular subtype, racial ancestry and clinical outcomes in prostate cancer. Modern Pathology 2018

32. Markowski MC, Hubbard GK, Hicks JL, Zheng Q, King A, Esopi D, et al. Characterization of novel cell lines derived from a MYC-driven murine model of lethal metastatic adenocarcinoma of the prostate. Prostate 2018;78:992–1000

33. Holdhoff M, Guner G, Rodriguez FJ, Hicks JL, Zheng Q, Forman MS, et al. Absence of cytomegalovirus in glioblastoma and other high-grade gliomas by real-time PCR, immunohistochemistry, and in situ hybridization. Clinical cancer research : an official journal of the American Association for Cancer Research 2017;23:3150–7

34. Doench JG, Fusi N, Sullender M, Hegde M, Vaimberg EW, Donovan KF, et al. Optimized sgRNA design to maximize activity and minimize off-target effects of CRISPR-Cas9. Nat Biotechnol 2016;34:184–91

35. Shalem O, Sanjana NE, Hartenian E, Shi X, Scott DA, Mikkelsen TS, et al. Genome-Scale CRISPR-Cas9 Knockout Screening in Human Cells. Science (New York, NY) 2014;343:84–7

36. Sanjana NE, Shalem O, Zhang F. Improved vectors and genome-wide libraries for CRISPR screening. Nat Methods 2014;11:783–4

37. Baena Del Valle J, Zheng Q, Hicks J, Fedor HL, Trock BJ, Morrissey C, et al. Rapid loss of RNA detection by in situ hybridization in stored tissue blocks and preservation by cold storage of unstained slides. American Journal of Clinical Pathology 2017;148:398–415

38. Kaur HB, Lu J, Guedes LB, Maldonado L, Reitz L, Barber JR, et al. TP53 missense mutation is associated with increased tumor-infiltrating T cells in primary prostate cancer. Hum Pathol 2019;87:95–102

39. Barretina J, Caponigro G, Stransky N, Venkatesan K, Margolin AA, Kim S, et al. The Cancer Cell Line Encyclopedia enables predictive modelling of anticancer drug sensitivity. Nature 2012;483:603–307

40. Hempel HA, Cuka NS, Kulac I, Barber JR, Cornish TC, Platz EA, et al. Low Intratumoral Mast Cells Are Associated With a Higher Risk of Prostate Cancer Recurrence. Prostate 2017;77:412–24

41. Baena-Del Valle JA, Zheng Q, Esopi DM, Rubenstein M, Hubbard GK, Moncaliano MC, et al. MYC drives overexpression of telomerase RNA (hTR/TERC) in prostate cancer. The Journal of Pathology 2018;244:11–24

42. Tomlins SA, Mehra R, Rhodes DR, Cao X, Wang L, Dhanasekaran SM, et al. Integrative molecular concept modeling of prostate cancer progression. Nat Genet 2007;39:41–51

43. Lapointe J, Li C, Higgins JP, van de Rijn M, Bair E, Montgomery K, et al. Gene expression profiling identifies clinically relevant subtypes of prostate cancer. Proceedings of the National Academy of Sciences of the United States of America 2004;101:811–6

44. Vanaja DK, Cheville JC, Iturria SJ, Young CY. Transcriptional silencing of zinc finger protein 185 identified by expression profiling is associated with prostate cancer progression. Cancer Res 2003;63:3877–82

45. Welsh JB, Sapinoso LM, Su AI, Kern SG, Wang-Rodriguez J, Moskaluk CA, et al. Analysis of gene expression identifies candidate markers and pharmacological targets in prostate cancer. Cancer Res 2001;61:5974–8

46. Wallace TA, Prueitt RL, Yi M, Howe TM, Gillespie JW, Yfantis HG, et al. Tumor immunobiological differences in prostate cancer between African-American and European-American men. Cancer Res 2008;68:927–36

47. Rhodes DR, Yu J, Shanker K, Deshpande N, Varambally R, Ghosh D, et al. ONCOMINE: a cancer microarray database and integrated data-mining platform. Neoplasia (New York, NY) 2004;6:1–6

48. Díez-Villanueva A, Mallona I, Peinado MA. Wanderer, an interactive viewer to explore DNA methylation and gene expression data in human cancer. Epigenetics & Chromatin 2015;8:22

49. Ahearn TU, Pettersson A, Ebot EM, Gerke T, Graff RE, Morais CL, et al. A Prospective Investigation of PTEN Loss and ERG Expression in Lethal Prostate Cancer. J Natl Cancer Inst 2016;108

50. Robinson D, Van Allen EM, Wu YM, Schultz N, Lonigro RJ, Mosquera JM, et al. Integrative clinical genomics of advanced prostate cancer. Cell 2015;161:1215–28

51. Haney NM, Faisal FA, Lu J, Guedes LB, Reuter VE, Scher HI, et al. PTEN Loss with ERG Negative Status is Associated with Lethal Disease after Radical Prostatectomy. J Urol 2020;203:344–50

52. Long T, He W, Pan Q, Zhang S, Zhang Y, Liu C, et al. Microglia P2X4 receptor contributes to central sensitization following recurrent nitroglycerin stimulation. Journal of Neuroinflammation 2018;15:245

53. Stokes L, Layhadi JA, Bibic L, Dhuna K, Fountain SJ. P2X4 Receptor Function in the Nervous System and Current Breakthroughs in Pharmacology. Front Pharmacol 2017;8

54. Matsumura Y, Yamashita T, Sasaki A, Nakata E, Kohno K, Masuda T, et al. A novel P2X4 receptor-selective antagonist produces anti-allodynic effect in a mouse model of herpetic pain. Scientific Reports 2016;6:32461

55. Chemicar N.

56. Suurväli J, Boudinot P, Kanellopoulos J, Rüütel Boudinot S. P2X4: A fast and sensitive purinergic receptor. Biomedical journal 2017;40:245–56

57. Srivastava P, Cronin CG, Scranton VL, Jacobson KA, Liang BT, Verma R. Neuroprotective and neuro-rehabilitative effects of acute purinergic receptor P2X4 (P2X4R) blockade after ischemic stroke. Experimental Neurology 2020;329:113308

58. Zumerle S, Calì B, Munari F, Angioni R, Di Virgilio F, Molon B, et al. Intercellular Calcium Signaling Induced by ATP Potentiates Macrophage Phagocytosis. Cell reports 2019;27:1-10.e4

59. Csóka B, Németh ZH, Szabó I, Davies DL, Varga ZV, Pálóczi J, et al. Macrophage P2X4 receptors augment bacterial killing and protect against sepsis. JCI Insight 2018;3

60. Erlandsson A, Carlsson J, Lundholm M, Falt A, Andersson SO, Andren O, et al. M2 macrophages and regulatory T cells in lethal prostate cancer. Prostate 2019;79:363–9

61. Zarif JC, Baena-Del Valle JA, Hicks JL, Heaphy CM, Vidal I, Luo J, et al. Mannose Receptor-positive Macrophage Infiltration Correlates with Prostate Cancer Onset and Metastatic Castration-resistant Disease. European urology oncology 2019;2:429–36

62. Ghalali A, Ye Z-w, Högberg J, Stenius U. PTEN and PHLPP crosstalk in cancer cells and in TGFβ-activated stem cells. Biomedicine & Pharmacotherapy 2020;127:110112

63. Mulholland DJ, Kobayashi N, Ruscetti M, Zhi A, Tran LM, Huang J, et al. PTEN Loss and RAS/MAPK Activation Cooperate to Promote EMT and Metastasis Initiated from Prostate Cancer Stem/Progenitor Cells. Cancer Research 2012;72:1878

64. Tomlins SA, Rhodes DR, Perner S, Dhanasekaran SM, Mehra R, Sun XW, et al. Recurrent fusion of TMPRSS2 and ETS transcription factor genes in prostate cancer. Science 2005;310:644–8

65. Tian TV, Tomavo N, Huot L, Flourens A, Bonnelye E, Flajollet S, et al. Identification of novel TMPRSS2:ERG mechanisms in prostate cancer metastasis: involvement of MMP9 and PLXNA2. Oncogene 2014;33:2204–14

