## Supplemental FIles for "P2X4 Purinergic Receptors as a Therapeutic Target in Aggressive Prostate Cancer"

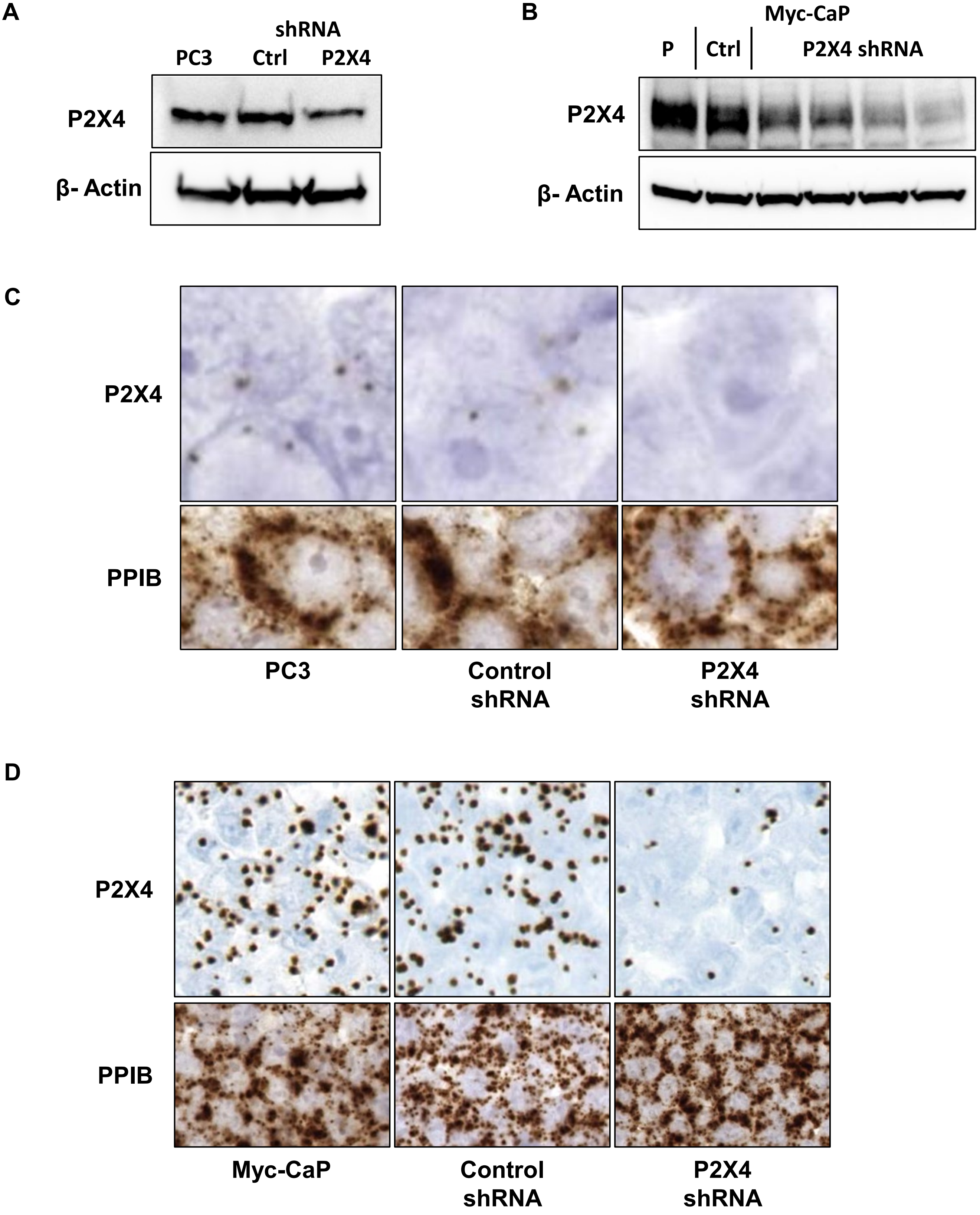


**Supplementary Figure 1**: P2X4 targeted shRNA shows specific reduction of (A) human anti-P2X4 antibody signal, (B) mouse anti-P2X4 antibody signal, (C) human P2X4 RISH probe signal and (D) mouse P2X4 RISH probe signal.

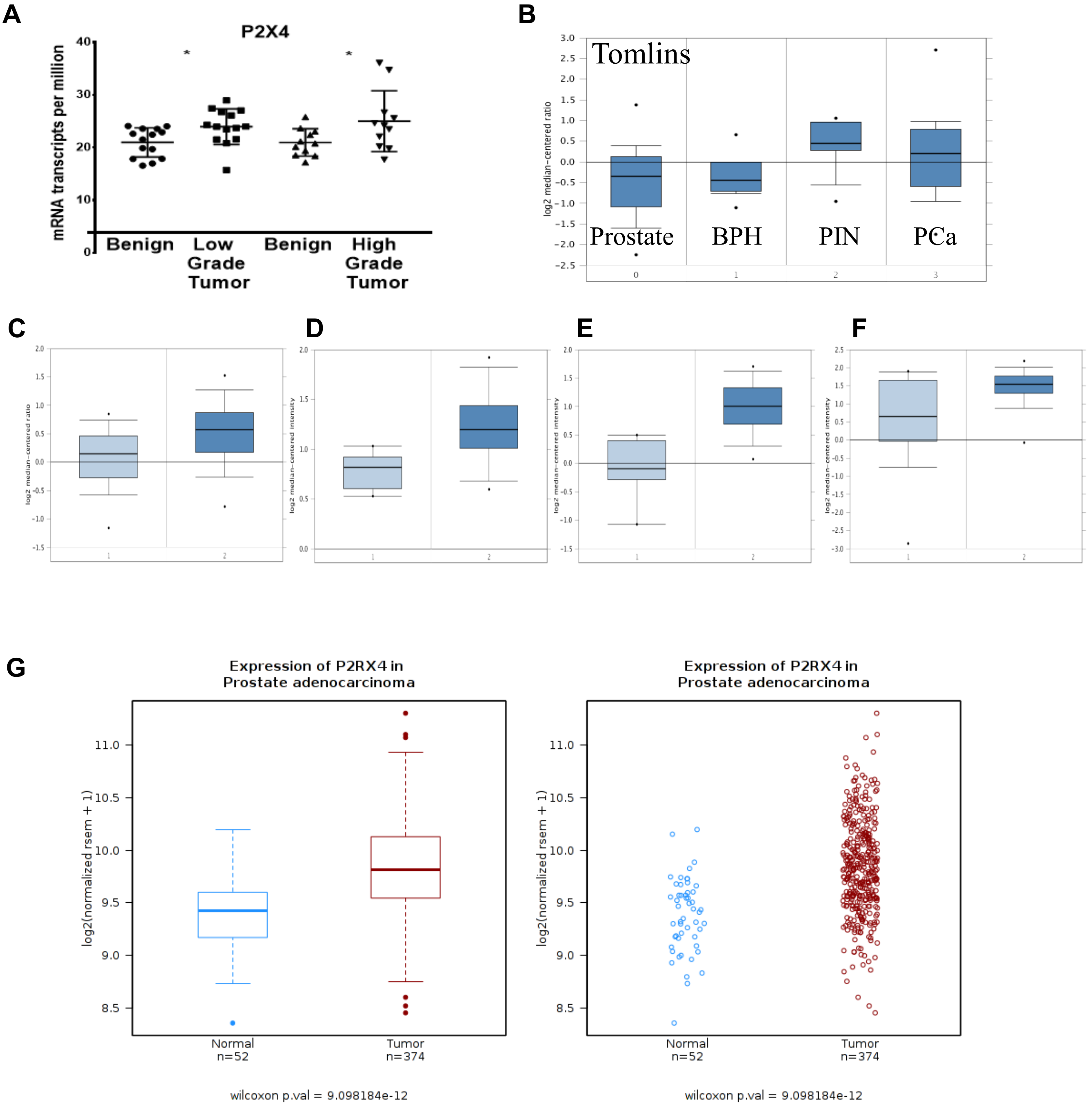


**Supplementary Figure 2**: (A) RNA sequencing analysis showed that P2X4 receptor expression is elevated in cancer compared to benign. Public gene datasets also show that P2X4 purinergic receptor mRNA expression is elevated in prostate cancer (dark blue) compared to benign prostate glands (light blue) in (B) Tomlins (p = 7.44E-5), (C)Lapointe (p = 8.81E-5), (D) Vanaja (p = 4.07E-5), (E)Welsh (p = 1.93E-5), and (F)Wallace (p = 0.002) datasets via ONCOMINE. Prostatic intraepithelial neoplasia (PIN) tissues also had elevated P2X4 mRNA expression compared to benign prostate in the Tomlins dataset. (G) There was significantly increased P2X4 mRNA expression in cancer compared to benign tissues in the TCGA dataset as assessed by Wanderer.

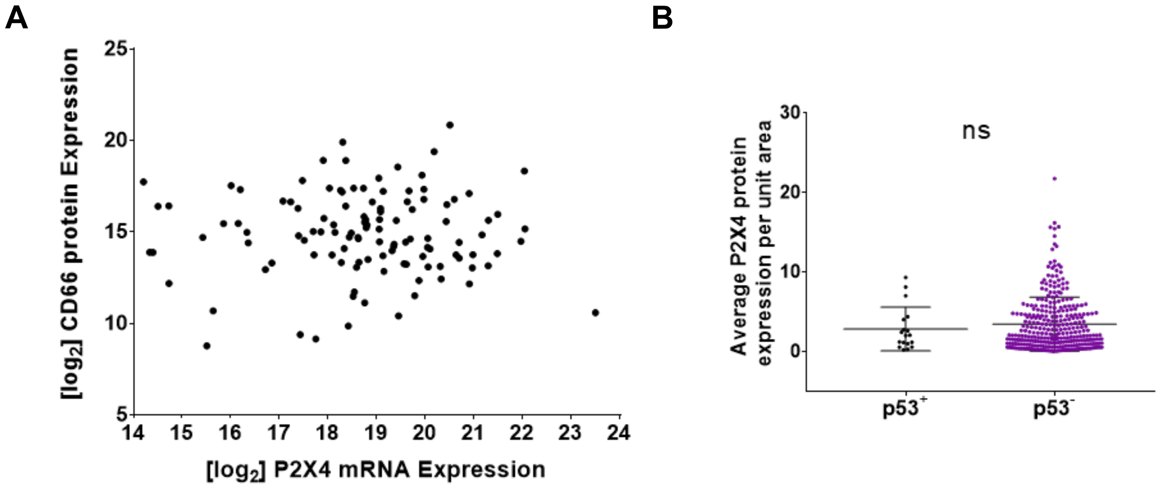


**Supplementary Figure 3**: (A) There was no correlation between P2X4 mRNA and CD66 protein expression in the PCBN High Grade Race TMA set. (B) There was no difference in P2X4 expression in cases with or without p53.

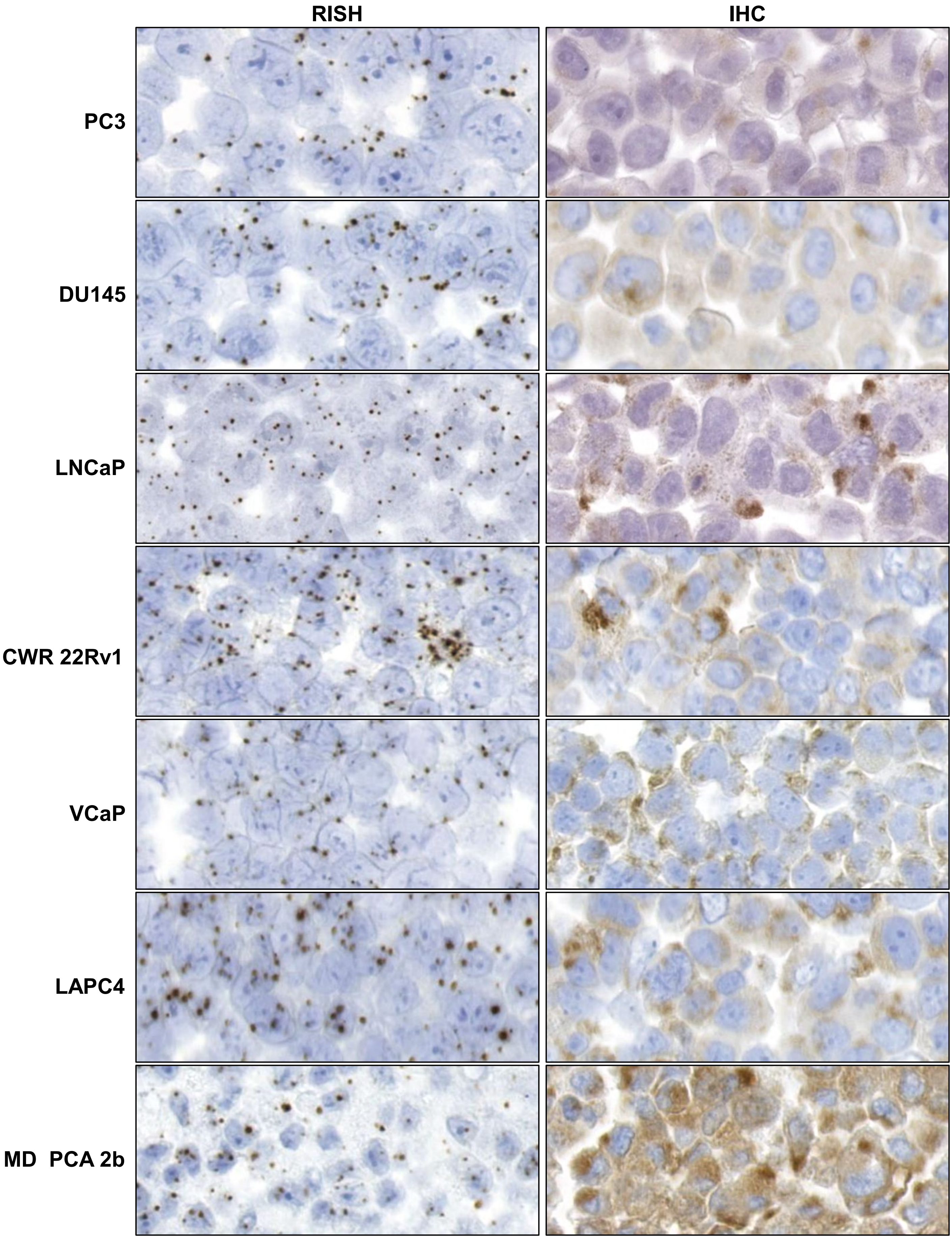


**Supplementary Figure 4**: P2X4 purinergic receptor mRNA and protein expression is detected in human derived prostate cancer cell lines

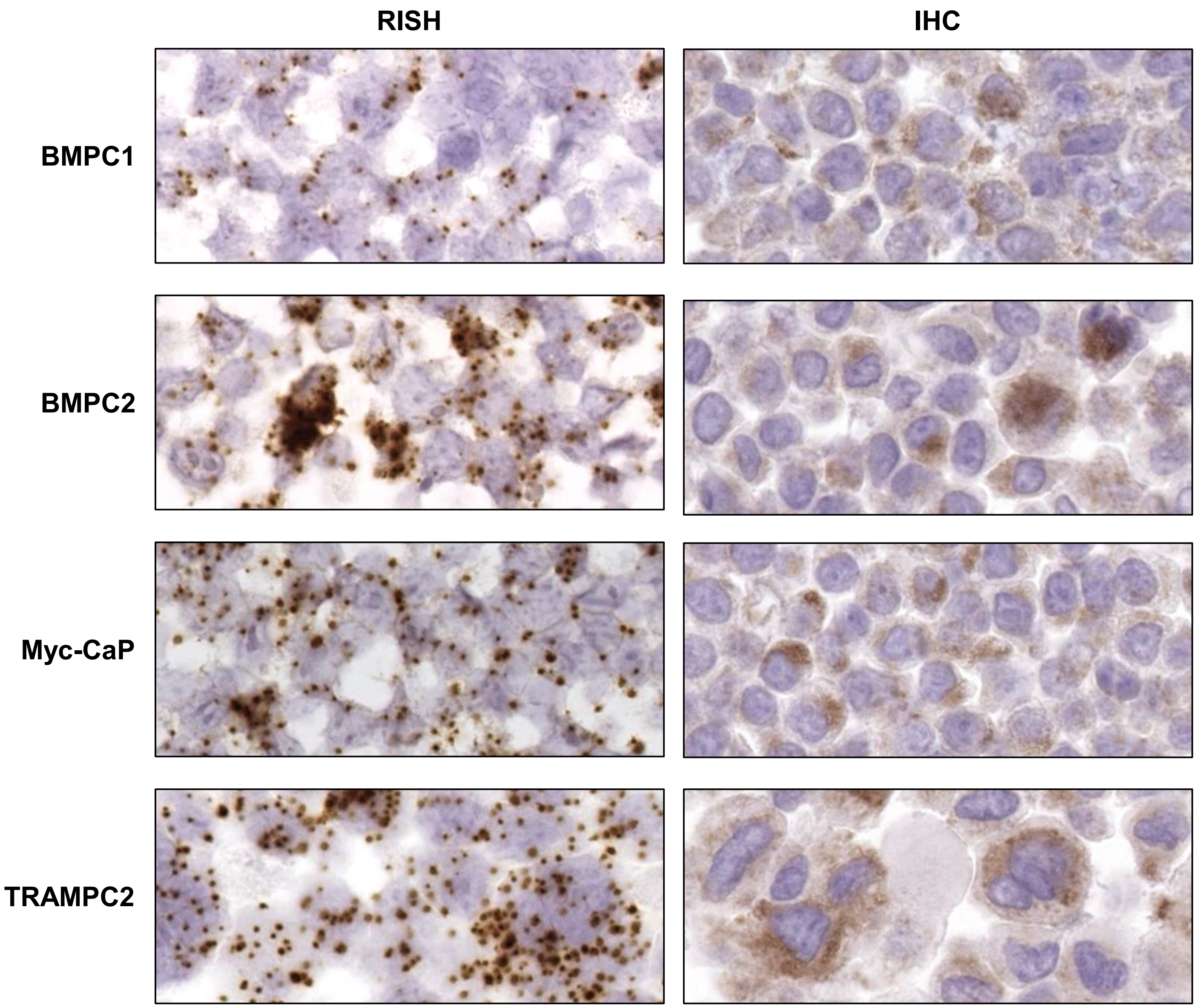


**Supplementary Figure 5**: P2X4 purinergic receptor mRNA and  protein expression is detected in mouse derived prostate cancer cell lines

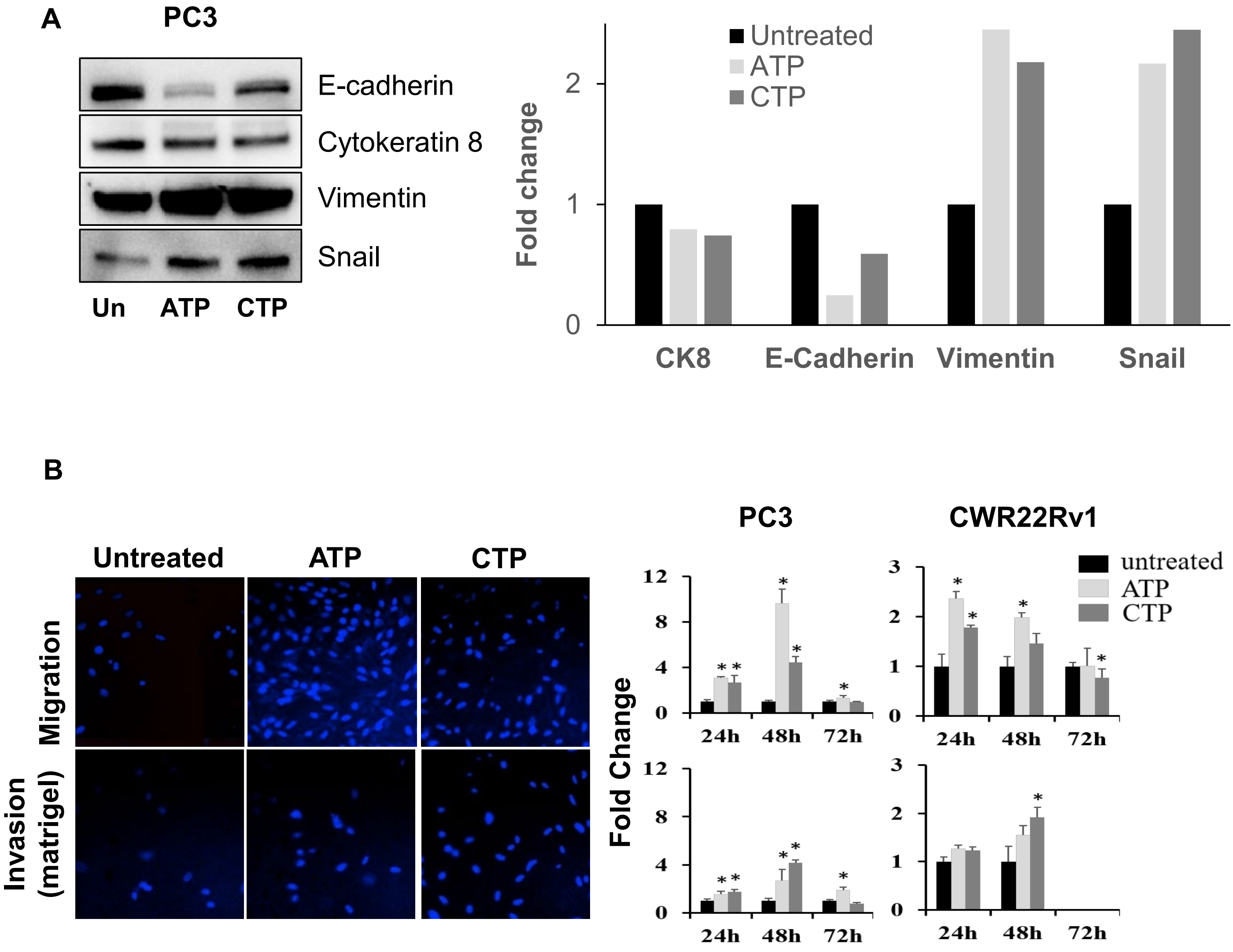


**Supplementary Figure 6:** Nucleotide treatment (A) decreased epithelial markers and increased mesenchymal markers and (B) increased migration and invasion of PCa cells.

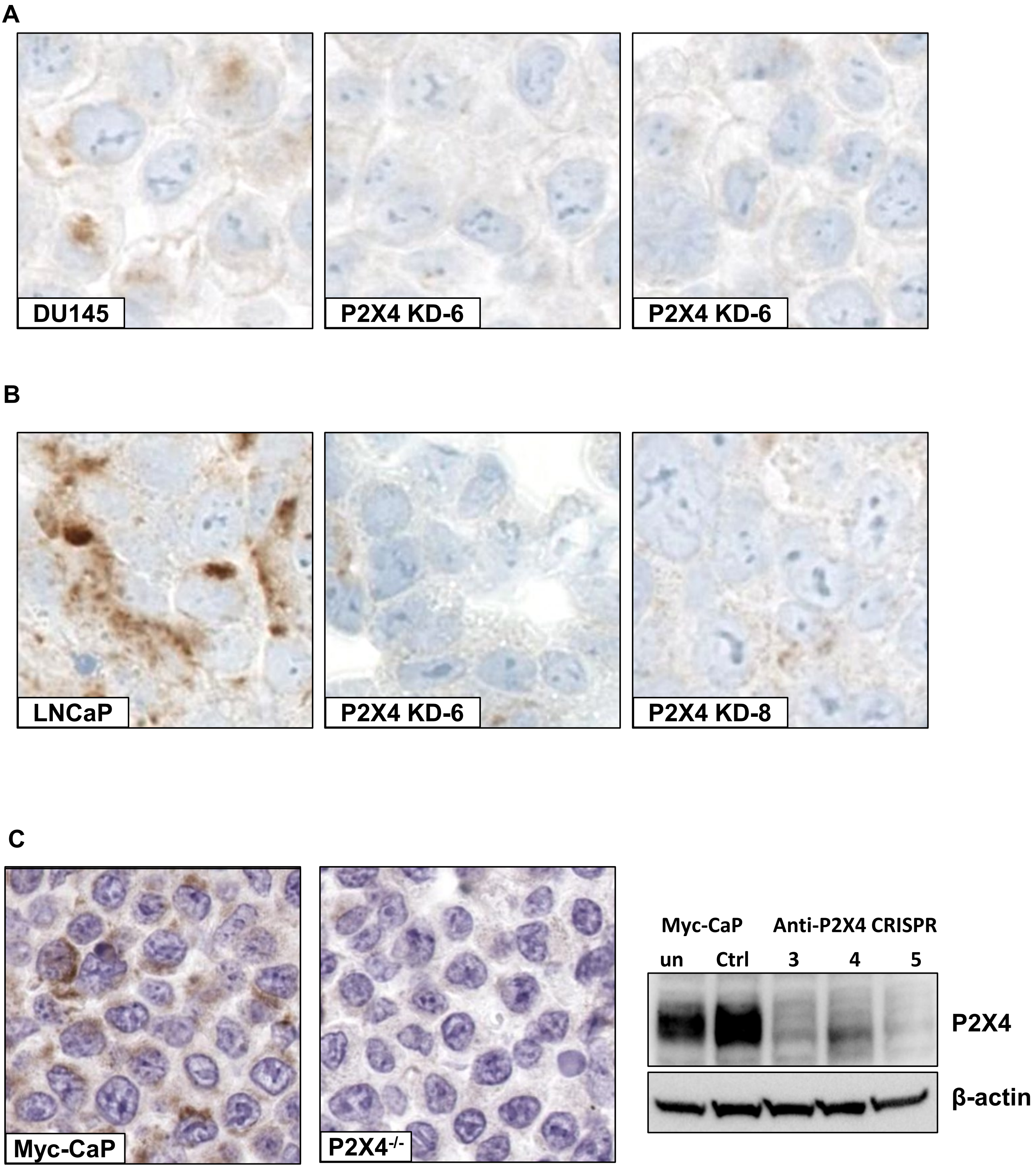


**Supplementary Figure 7:** CRISPR knockdown of P2X4 is confirmed by RISH analysis of (A) DU145, (B) LNCaP and (C) Myc-CaP cells. Anti-P2X4 CRISPR 5, shown by western blot, was selected for inoculation into mice.

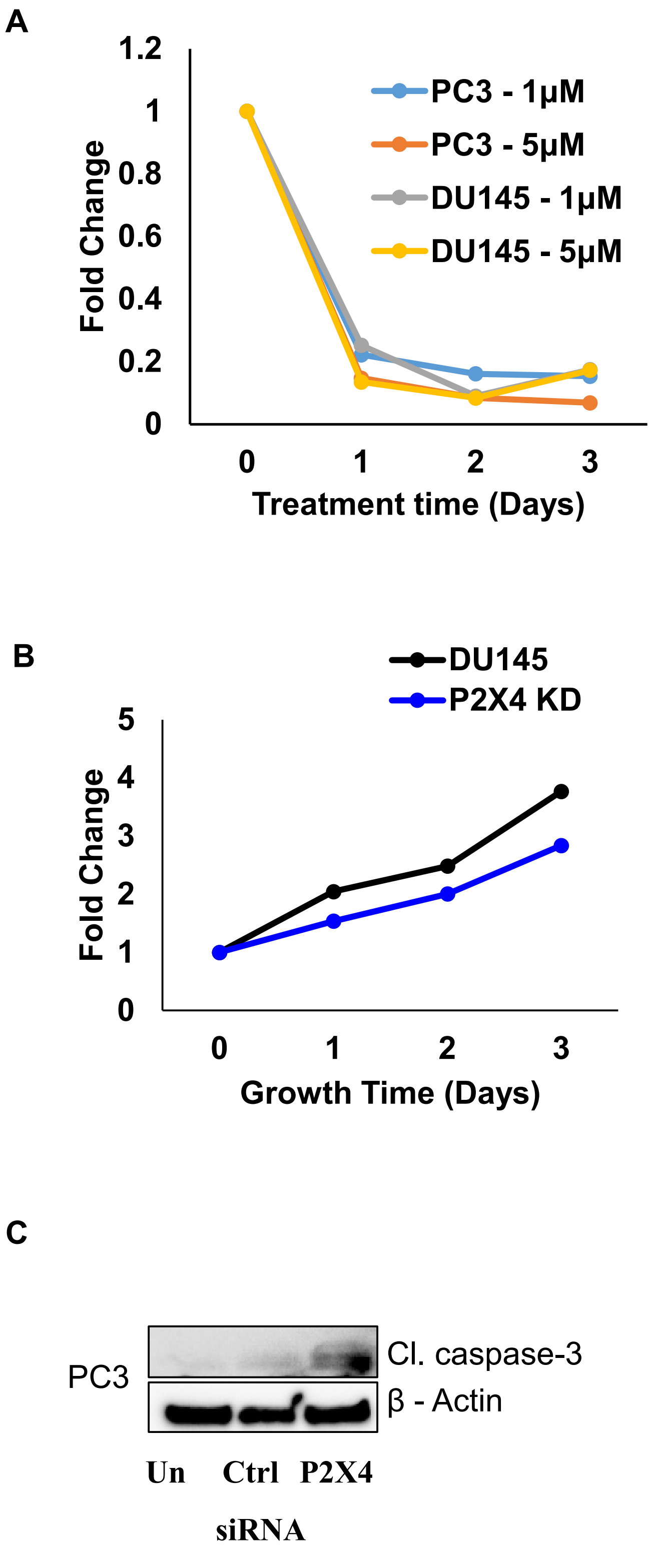


**Supplementary Figure 8**: (A) Treatment of PC3 cells with 5-BDBD decreased cell viability at 24, 48 and 72h. (B) P2X4 knockdown reduced DU145 cell viability. (C) P2X4 siRNA temporary knockdown induced cleaved caspase-3 protein expression.

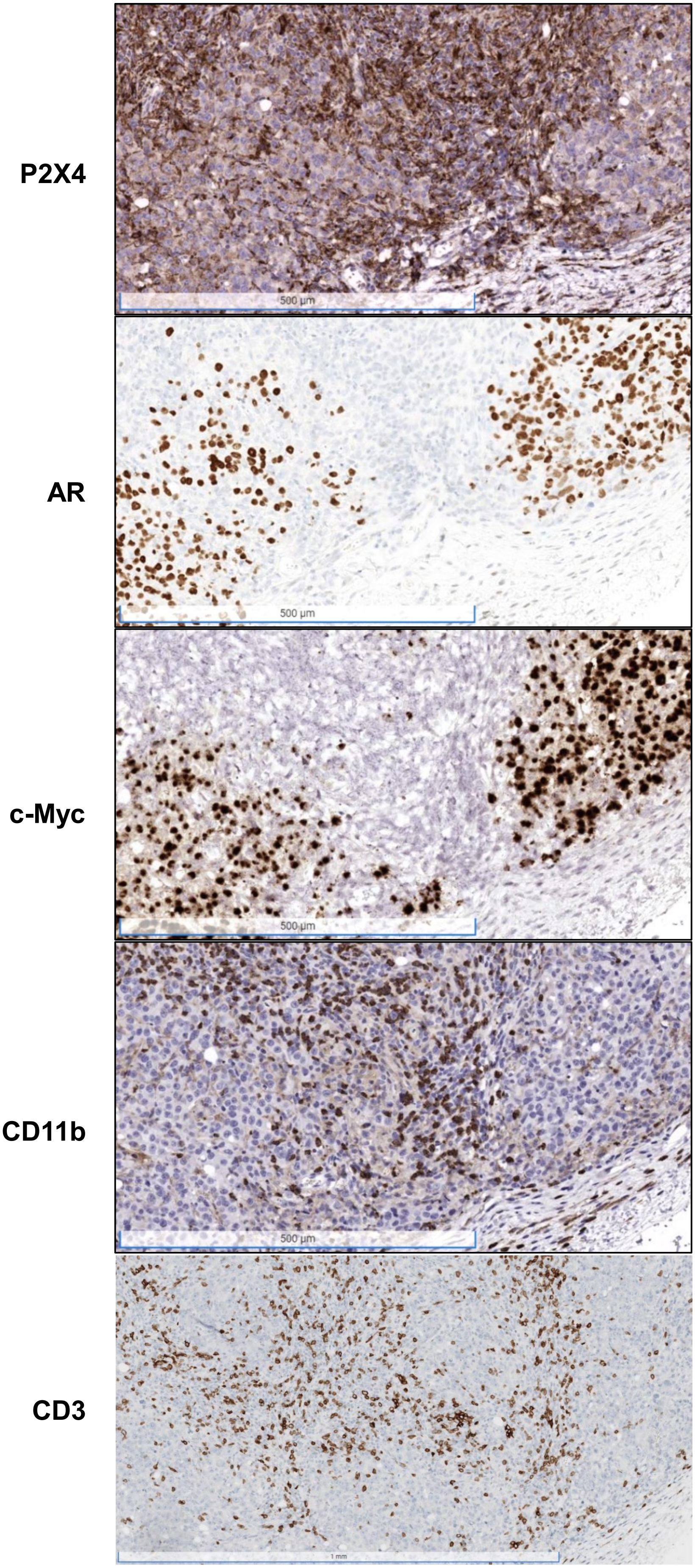


**Supplementary Figure 9** P2X4 positive regions lacking AR and human c-Myc stain positive for CD11b in Myc-CaP allograft tissue from mice

**Supplementary Table S1**. Patient characteristics for whole tissue sections.

| **Organ Donor Prostates** | Black | White |
|---|---|---|
| Median Age, Years | 21 | 23 |
| Race, # cases | 1 | 3 |

| **Whole Tissue Sections** | Black | White |
|---|---|---|
| Median Age, Years | 58 | 60 |
| Race, # cases | 3 | 3 |
| Gleason, # cases | | |
| <=6 | 2 | 2 |
| 3+4 | 1 | 1 |
| 4+3 | 0 | 0 |
| >=8 | 0 | 0 |
| Stage, # cases | | |
| T2 | 2 | 2 |
| T3 or Above | 1 | 1 |

**Supplementary Table S2**. Patient characteristics for TMA sets.

| **Race Disparity Case TMA Set** | Black | White | Other |
|---|---|---|---|
| Median Age, Years | 58 | 60 | NA |
| Race, # cases | 139 | 151 | 1 |
| Gleason, # cases | | | |
| <=6 | 33 | 32 | 0 |
| 3+4 | 41 | 42 | 0 |
| 4+3 | 32 | 38 | 1 |
| >=8 | 33 | 39 | 0 |
| Stage, # cases | | | |
| T2 | 54 | 72 | NA |
| T3 or Above | 67 | 79 | NA |
| Unknown | 18 | 0 | NA |

| **PCBN High Grade Race TMA Set** | Black | White | Other |
|---|---|---|---|
| Median Age, Years | 61 | 62 | NA |
| Race, # cases | 120 | 60 | 0 |
| Gleason, # cases | | | |
| <=6 | 20 | 10 | |
| 3+4 | 26 | 13 | |
| 4+3 | 20 | 10 | |
| >=8 | 54 | 27 | |
| Stage, # cases | | | |
| T2 | 73 | 36 | |
| T3 or Above | 47 | 24 | |

| **Metastasis at Autopsy TMA Set** | Black | White | Other |
|---|---|---|---|
| Median Age, Years | 66 | 70 | |
| Race, # cases | 5 | 16 | 0 |
| Sites, # sampled | 4 | 8 | |
| # spots | | | |
| Prostate benign | 6 | 9 | |
| Prostate cancer | | | |
| Seminal vesicle | 0 | 5 | |
| Liver | 6 | 9 | |
| Lung | 6 | 5 | |
| Lymph Node | 12 | 53 | |
| Bone | 15 | 86 | |
| Adrenal Gland | 0 | 3 | |
| Skin | 0 | 6 | |
| Spleen | 0 | 3 | |
| Bladder | 0 | 2 | |
| Pancreas | 0 | 1 | |
| Other soft tissue | 0 | 4 | |

**Supplementary Table S3**. Association between P2X4 expression and prostate cancer metastasis, or biochemical recurrence by race.*

| Metastasis | Case (N) | Control(N) | HR (95% CI) ** | | | *p* |
|---|---|---|---|---|---|---|
| **White Patients** | | | | | | |
| *Cancer* | | | | | | |
| Below Median | 6 | 64 | | Ref | | |
| Above Median | 15 | 67 | 1.617 | 0.592 | 4.415 | 0.3484 |
| *Benign* | | | | | | |
| Below Median | 3 | 68 | | Ref | | |
| Above Median | 18 | 63 | 4.362 | 1.252 | 15.199 | 0.0207 |
| **Black Patients** | | | | | | |
| *Cancer* | | | | | | |
| Below Median | 6 | 61 | | Ref | | |
| Above Median | 8 | 47 | 1.091 | 0.205 | 5.811 | 0.919 |
| *Benign* | | | | | | |
| Below Median | 7 | 59 | | Ref | | |
| Above Median | 7 | 49 | 1.196 | 0.33 | 4.333 | 0.7849 |
| Biochemical Recurrence | | | | | | |
| **White Patients** | | | | | | |
| *Cancer* | | | | | | |
| Below Median | 26 | 44 | | Ref | | |
| Above Median | 31 | 51 | 1.02 | 0.595 | 1.746 | 0.9433 |
| *Benign* | | | | | | |
| Below Median | 20 | 51 | | Ref | | |
| Above Median | 37 | 44 | 1.552 | 0.889 | 2.708 | 0.122 |
| **Black Patients** | | | | | | |
| *Cancer* | | | | | | |
| Below Median | 21 | 46 | | Ref | | |
| Above Median | 23 | 32 | 1.634 | 0.806 | 3.315 | 0.1736 |
| *Benign* | | | | | | |
| Below Median | 20 | 46 | | Ref | | |
| Above Median | 24 | 32 | 1.298 | 0.703 | 2.397 | 0.4051 |

*, participants from Race Disparity" and "PCBN High Grade Race" TMA sets. Median was determined based on the P2X4 measurement among all men. **, hazard ratio (HR) and 95% confidence intervals (CI) estimated from Cox Hazard regression model adjusting for age, stage, grade, TMA sets and PTEN status.
